# Multi-step antibody class switching in a primary human response is restricted after *IGHG2* and dependent on B cell maturation stage

**DOI:** 10.1101/2025.05.07.652638

**Authors:** Guillem Montamat Garcia, Joseph C. F. Ng, Alexander T. Stewart, Emma Sinclair, Paul Blair, Diana Kateregga, Amir Gander, David Kipling, Dongjun Guo, Lutecia Servius, Christopher J. M. Piper, Zara Baig, Franca Fraternali, Claudia Mauri, Deborah K. Dunn-Walters

## Abstract

Class switch recombination (CSR) allows the formation of functionally specialized antibodies. Understanding of CSR dynamics is key for better design and prediction of vaccines to protect mucosal surfaces. To investigate CSR in a primary human immune response under a controlled setting, we sampled healthy volunteers without COVID-19 history every other day during the first three weeks after SARS-CoV-2 vaccination, with additional time points up to six months. Leveraging bulk and single-cell B cell receptor repertoires, single-cell transcriptomics, and immunophenotyping, we uncover paradigm-shifting insights into CSR. Newly activated B cells produce sterile transcripts of all antibody isotype constant regions (*IGHC*) simultaneously up to *IGHG2*, challenging the view that sterile transcription occurs for only a single *IGHC* gene at a time. CSR follows a multistep progression along the *IGHC* locus; in this challenge vaccine-induced B cells switch from *IGHM* to *IGHG3* and *IGHG1*, followed by a subsequent switching to *IGHA1* and *IGHG2* after secondary immunization. *IGHA2* clones require pre-switching to *IGHA1* clones. Notably, switching tendency, measured by *IGHC* sterile transcription, is memory B cell subtype dependent, particularly beyond *IGHG2*. Contrary to other vaccines, antigen-specific B cells are enriched in DN2, Cmem2 and DN4 subtypes after immunization. We also observe a temporal decoupling of CSR and somatic hypermutation (SHM), with the latter detectable only after six months post-immunization. Our data describes the dynamics between CSR, SHM, sterile transcription and B cell memory development during a human primary response, challenges textbook models of CSR and offers new insights to aid control of CSR direction.

**Graphical Abstract:** 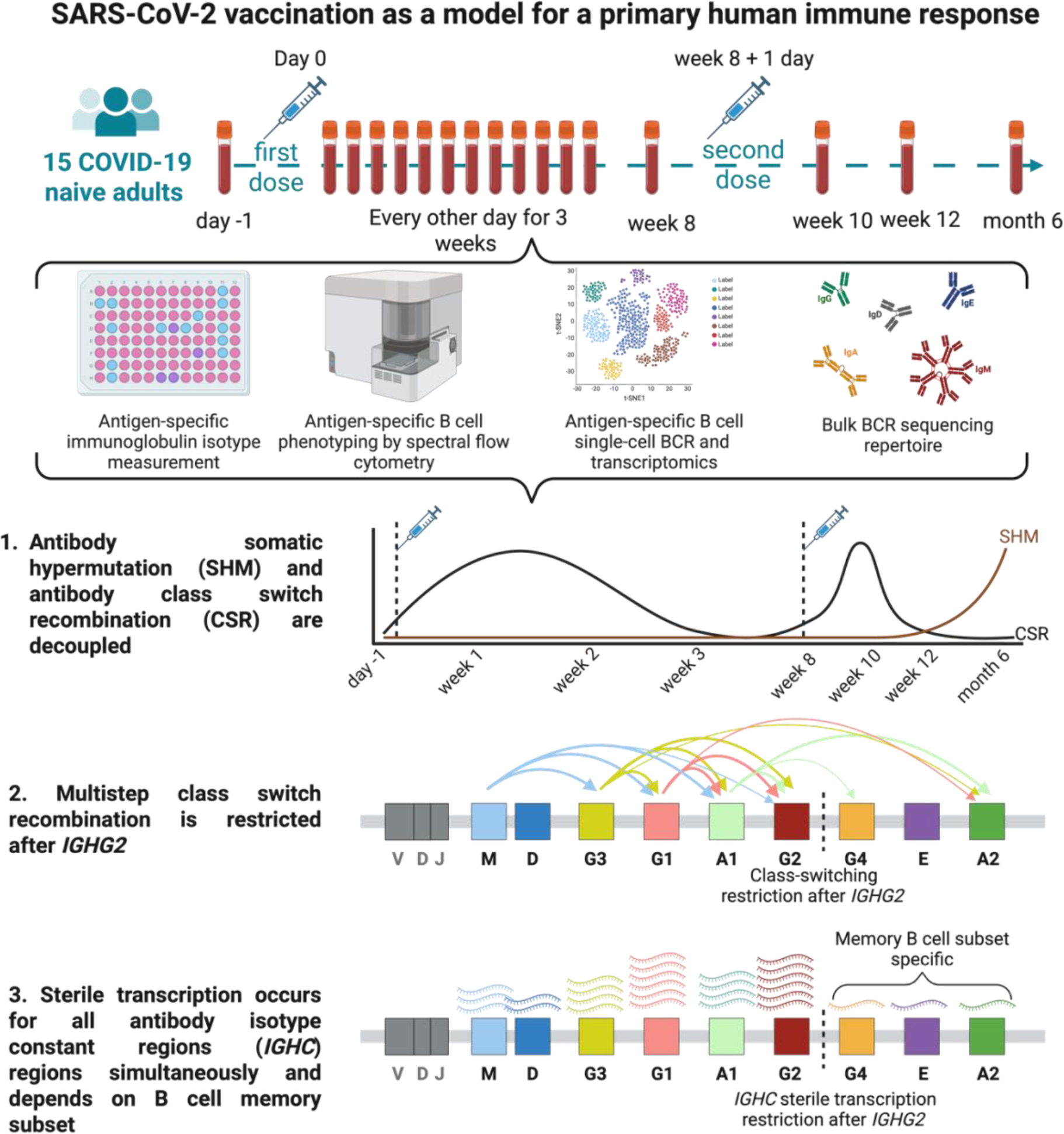

**One Sentence Summary:** Class switch recombination occurs independently of somatic hypermutation and in a multistep fashion up to *IGHG2* during a primary response in humans.

## Introduction

B cells, the mainstay of humoral responses, exist in many different developmental and functional forms, including regulators, antigen presenters, T cell interactors as well as antibody producers. Since B cells are very often underrepresented and poorly analyzed in immune single cells RNA sequencing (scRNA-seq) data, public cell atlases are lacking in crucial B cell information, especially during primary immune responses. B cells confer specificity against immune challenges via immunoglobulins (Ig) which can be either membrane bound (B cell receptor or BCR) or secreted in the form of antibodies(*1*). In humans, there are 5 isotypes, or classes, of B cell receptors (BCRs) with subtypes for IgG and IgA (IgM, IgD, IgE, IgG1/2/3/4, IgA1/2)(*2*). Identifying specific BCR isotypes is important as they are linked to distinct immune functions and responses. For example, IgA is predominantly localized to mucosal surfaces, including those in the respiratory and gastrointestinal tracts(*3–5*), while IgG, the most abundant antibody isotype in serum, plays a key role in immune responses against viral infections(*6–8*). The subtype of a BCR can also show important effector and functional differences, i.e. IgG1 and IgG2 have different Fc receptor binding affinities(*9, 10*), IgA2 can dimerize and be secreted across mucosal surfaces while IgA1 remains mainly systemic (*11*). Distinguishing between subtypes in high-resolution, single-cell settings has been difficult in immunology, but newer genomic and transcriptomic techniques can provide information on the type of B cell(*12*), its maturity in a response(*13*) as well as fundamental B cell processes such as class-switch recombination (CSR)(*14*). This new information can provide key insights into the role of B cells in disease conditions such as autoimmunity or responses to antigenic challenges of infection and vaccination(*15–18*).

B cells switch their BCR subtypes via class-switch recombination (CSR), where B cells undergo irreversible DNA recombination at their immunoglobulin heavy-chain (*IGH*) loci(*19–21*) to bring a different constant region gene (*IGHC*) into proximity with the variable region to form a productive immunoglobulin (Ig) transcript. The irreversibility of these events, due to genomic deletion, imposes a 5’ to 3’ directionality on CSR(*19, 20*). The use of transcription to open the genomic DNA to cutting and splicing factors during CSR results in the production of sterile *IGHC* transcripts. These sterile transcripts can be detected in scRNA-seq data and serve as markers to predict the likely future directionality of CSR(*22*). Our recent development of dedicated computational pipelines(*14*) has facilitated the study of CSR directionality preferences. However, a knowledge gap exists regarding the role of human B cell CSR during primary immune responses, particularly in understanding the dynamics of CSR and potential preferential pathways to specific final *IGHC* when naive individuals encounter an antigen for the first time. It has long been debated whether CSR occurs directly (exclusively from IgM-expressing naive B cells), indirectly (via intermediate *IGHC* genes) or a combination of the two(*19, 20, 23*). Most adult human immune response studies on vaccinated individuals are limited in longitudinal follow up and involve vaccine challenges within the geography of endemic disease, which means they will likely be a secondary challenge to a previous primary exposure. Although animal models allowed the characterization of CSR in a primary immune response(*24*), there are substantial differences in the genomic architecture of the *IGHC* locus between mice and humans(*25–27*).

Here we took advantage of an antigen-naive cohort being vaccinated with the SARS-CoV-2 mRNA-1273 vaccine to obtain detailed multi-omic B cell data from samples taken at frequent time points post-vaccination. This primary response data includes bulk and single-cell BCR sequencing, single-cell transcriptomics, and flow cytometry immune phenotyping of whole blood and isolated lymphocytes along with saliva samples collected under a post-vaccination (1^st^ and 2^nd^ doses) strict schedule, as well as a 6 months post-immunization timepoint. This allowed us to finely map B cell and CSR dynamics during primary immunization and study vaccine-derived antigen-specific B cells. Data of this multi-omic resource can be visualized in an integrated and interactive web browser (https://fraternalilab.cs.ucl.ac.uk/CovVaxBcells/).

We reveal that CSR follows a multistep pattern with a predominant preference for "vicinity switching" to adjacent *IGHC.* Activated B cells produce sterile transcripts of all *IGHC* regions up to *IGHG2,* with restricted transcription and class switching beyond the *IGHG2* locus. This changes the textbook understanding of CSR dynamics and is important knowledge for researchers to manipulate CSR for specific outcomes e.g. mucosal antibody generation in vaccination. We also observed the emergence of antigen-specific atypical B cell subsets, such as double negative (DN) 2 B cells, particularly following the second vaccine dose. Despite an early high class-switching rate, a lack of somatic hypermutation (SHM) was observed in antigen-specific B cells until 6 months post-immunization, which is in contrast with the classical view where CSR and SHM occur simultaneously in the germinal center (GC). Indeed, our novel results in humans, concur with evidence in mice showing that CSR and SHM are decoupled(*28*). This knowledge will need to be incorporated into our future thinking around the dynamics of human responses. Furthermore, our study provides comparable bulk versus antigen-specific BCR repertoires, enabling the testing of prior assumptions in the field that all expanded Ig genes in a response encode high affinity antibodies against the challenging antigen. This observed divergence between these paired data sets emphasizes the importance of multi-omic approaches for a nuanced understanding of antigen-specific responses. Overall, our data describe the dynamics of CSR, SHM, sterile transcription, and B cell memory development during a primary response in naive humans with unprecedented detail, challenging textbook and previous CSR models and offering new insights to help direct CSR control.

## Results

### Antigen-specific, class-switched, CD27-negative memory B cells appear after the second dose of SARS-CoV-2 mRNA vaccine

We took advantage of the SARS-CoV-2 mRNA-1273 vaccination as a model to address the knowledge gap in human CSR dynamics and B cell clonal evolution during primary responses. Fifteen healthy adults were immunized with the SARS-CoV-2 mRNA-1273 vaccine and samples collected for multi-omic measurement using a comprehensive schedule with timepoints every other day for the first three weeks (details in methods and **Fig. 1A**). Two (P6 and P14) out of fifteen participants were excluded after testing positive for anti-receptor binding domain (RBD) of the SARS-CoV-2 Spike (S) protein IgG antibodies, indicating prior SARS-CoV-2 infection (Supp. Fig. 1A).

**Figure 1.**
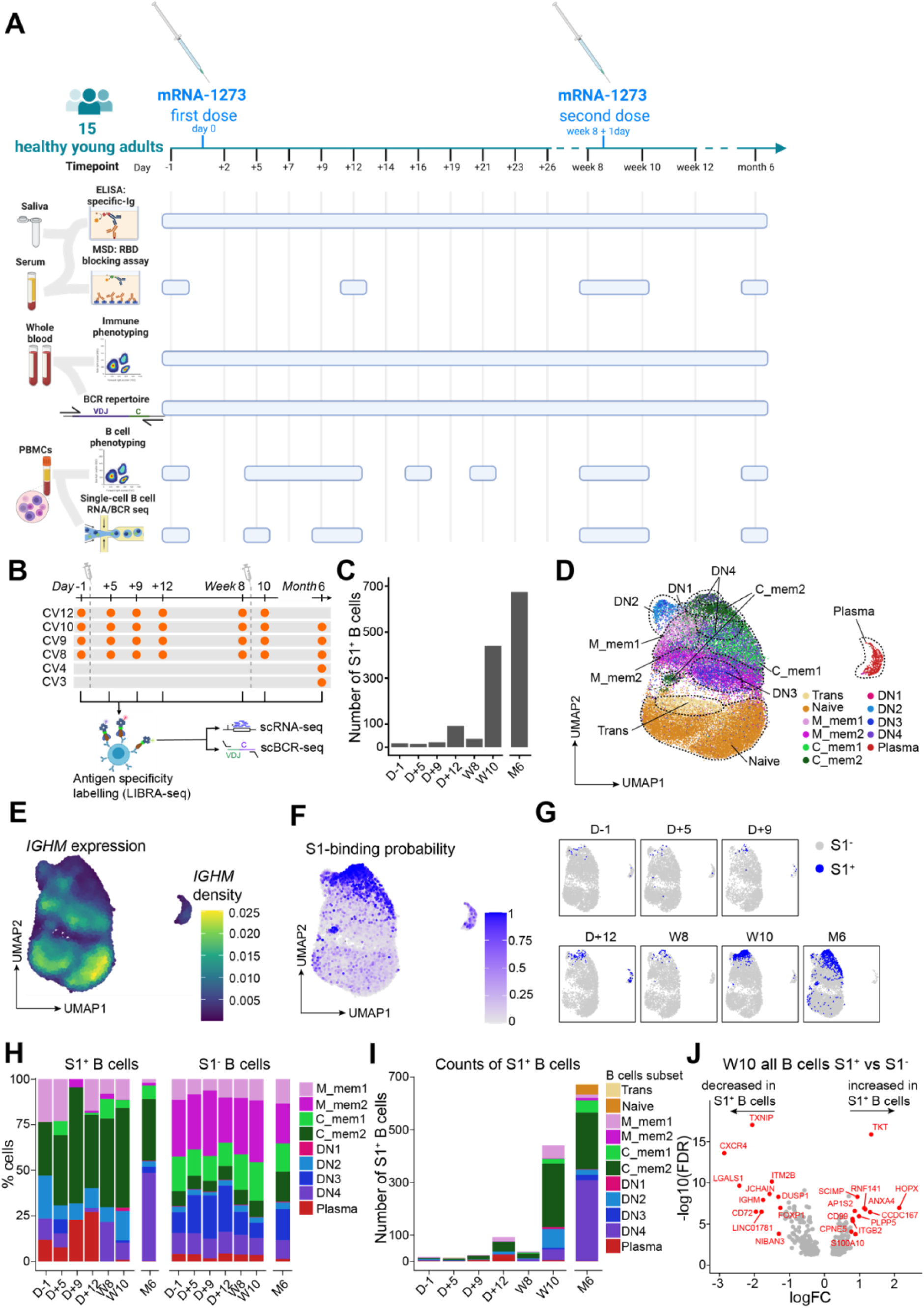
Multi-immunomic SARS-CoV-2 vaccination study detects antigen-specific class-switched double-negative (DN) memory B cells after the second dose. (A) Schematic illustrating sample collection timeline and assays performed on the collected samples. The schedule included a baseline on Day −1, with initial vaccination on Day 0, followed by time points every Monday, Wednesday, and Friday from Day 2 to Day 26. The second dose was administered at week 8, with pre-dose baseline (W8) and post-dose time points at Week 10 and Week 12, concluding with a final check at 6 months post-initial vaccination. (B) Schematic illustration of time points selected for single-cell transcriptomic profiling. (C) Number of S1^+^ B cells captured in single-cell transcriptomic data across time points. (D) Projection of scRNA-seq data of all B cells (S1^+^ and S1^-^ B cells) using Uniform Manifold Approximation and Projection (UMAP). Cell labels are transferred from a previously published scRNA-seq atlas of peripheral B cells based on gene expression similarities (see Methods for details). (E) *IGHM* gene expression per cell in the scRNA-seq data and visualized on the UMAP projection. (F) S1-binding probability score quantified per cell in the scRNA-seq data and visualized on the UMAP projection. (G) S1^+^ (blue) and S1^-^ (gray) B cells at each time point assayed in scRNA-seq visualized using the UMAP projection overtime. (H) Relative frequency distribution of S1^+^ and S1^-^ memory B cells and plasma cells across time. (I) Number of S1^+^ B cells sampled at each timepoint grouped by B cell subset labels. (J) Differential gene expression analysis between S1^+^ and S1^-^ B cells at the W10 time point across all B cell subsets.

Vaccine-induced RBD-specific IgG, IgA, and IgM antibodies emerged in the serum by day 12 post vaccination (D+12), with IgG increasing significantly after the second dose, while IgA and IgM remained unchanged; unlike in infection, salivary IgA did not rise during vaccination (Supp. Fig. 1B-C). By month 6 (M6), all RBD-specific antibodies declined, though sera blocking capacity against multiple SARS-CoV-2 strains persisted, except against Omicron variants, which showed an unexpected reduction compared to baseline (Supp. Fig. 1B,D-K). A rapid surge of antibody-secreting cells peaked at D+7 and D+9 without altering total B cell ratios (Supp. Fig. 2A-C), indicating targeted activation rather than global expansion, and highlighting the importance of early response assessments.

Vaccine-derived, Spike protein subunit 1 (S1)-specific (S1^+^) B cells from blood samples were labeled with a double-conjugated (nucleotide barcode and fluorochrome) S1 bait using the LIBRA-seq technology(*29*) and processed using flow-activated cell sorting (FACS) followed by 5’ single-cell transcriptomics (10X Chromium). This approach enabled us to generate matched single-cell RNA sequencing (scRNA-seq) and scBCR-seq datasets of the same cells, some of which were known to bind the antigen. Guided by the antibody (Supp. Fig. 1B), cellular (Supp. Fig. 2B) and repertoire (Supp Fig. 3) responses, we selected key timepoints for this analysis (**Fig. 1B**). In total, we analyzed the transcriptional profile of 27,027 B cells up to week 10 (W10), of which 623 were S1^+^ from four participants. Consistent with serum titers (Supp. Fig. 1B) and flow cytometry measurements (Supp. Fig. 2D-E), most S1^+^ B cells identified by single-cell transcriptomics were from W10, with smaller numbers observed at D+12 (**Fig. 1C**). A separate single-cell experiment was performed for month 6 (M6) samples with the aim to evaluate the long-term effects of the vaccine. A total of 7,308 S1^−^ B cells and 674 S1^+^ B cells were identified for this later timepoint from five participants (**Fig. 1B**).

Using a previously published transcriptomic atlas of healthy peripheral B cells(*12*) we identified 11 B cell clusters (**Fig. 1D**), each exhibiting distinct transcriptional signatures (Supp. Fig. 2F). When assessed for class-switch status, we were able to distinguish switched (*IGHM*-) and non-switched (*IGHM*+) B cells (**Fig. 1E**). Most S1^+^ B cells mapped onto the same uniform manifold approximation and projection (UMAP) space as switched cells, memory cells and the plasmablasts cluster (**Fig. 1E-G**), indicating the predominant class-switched status of S1^+^ B cells.

The antigen-specific B cell response was characterized by the expansion of C_mem2 B cells (a rare subtype in S1^-^ B cells), DN2 B cells and plasmablasts after primary immunization (D+12) (**Fig. 1H-I**). After the second vaccine dose (W10), S1^+^ C_mem2, DN2 and DN4 B cells increased in frequency (**Fig. 1H-I**). Although S1^+^ frequencies overall declined after six months (M6) (Supp. Fig. 2E) most S1^+^ B cell subsets were maintained, with the exception of DN2 which were reduced in favor of increased levels of DN4. (**Fig. 1H**). S1^+^ plasmablasts expanded predominantly at D+12, with no further increase after the second dose (**Fig. 1H-I**).

Flow cytometry phenotyping corroborated these dynamics, confirming S1^+^ plasmablast expansion at D+12 and the increase in S1^+^ DN B cells, particularly after the second dose (W10) (Supp. Fig. 4A-C). Additionally, the balance between unswitched and switched S1^+^ memory B cells shifted towards switched memory B cells only after the second dose (W10) (Supp. Fig 4B-C). In contrast, the phenotypes of S1^-^ B cells remained largely unchanged (**Fig. 1H**), apart from a reduction in both switched and unswitched memory B cells at D+9 and a two-fold increase in S1^-^ plasmablasts at D+7 (Supp. Fig. 4D).

Next, we analyzed the differences between S1^+^ and S1^-^ B cells at W10, the peak of the antigen-specific response (Supp. Fig. 2E). The transcriptional signature of S1^+^ B cells (**Fig. 1J**) suggested B cell activation and proliferation, including upregulation of *RNF141*, *CCDC167*, *HOPX* and downregulation of *IGHM*, *DUSP1*, *LINC01781, CD72* and *NIBAN3*. Additional differentially expressed pathways included B cell migration (*CD99* and *ITGB2*), MHC-II antigen presentation (*SCIMP* and *AP1S2*), NFkB signaling (*ANXA4*), calcium signaling (*CPNE5* and *S100A10*) and the germinal center response (upregulation of *PLPP5* and suppression of *ITM2B*, *CXCR4, FOXP1*). Metabolic pathways were also differentially expressed, including the pentose phosphate pathway (*TKT*) and redox pathways (*TXNIP*), the latter being the most downregulated gene in S1^+^ B cells. *JCHAIN,* associated with Blimp1-driven antibody-secreting cell (ASC) differentiation(*30*), was downregulated, suggesting a reduced propensity for S1^+^ B cells to become ASCs at W10. Notably, *LGALS1*, encoding galectin-1, a factor linked to B regulatory and plasma cell formation(*31*), was also reduced in S1^+^ B cells (**Fig. 1J**).

Within overrepresented subsets such as C_mem2, DN2, and DN4, S1^+^ cells exhibited transcriptional profiles distinct from their S1^-^ counterparts (Supp. Fig. 5), except for *TXNIP*, which was consistently downregulated across all subsets. These profiles reflected the activated state of S1^+^ B cells, aligning with prior studies(*32, 33*).

Taken together, our scRNA-seq resource identifies S1^+^ B cells as class-switched, C_mem2 and DN cells, with a distinct transcriptional signature. The induction of antigen-specific DN B cells, especially after the 2^nd^ dose was unexpected as this B cell subtype, particularly the DN2, has been associated with autoimmunity and aging(*34–37*). This unusual response during vaccination could be due, at least partially, to the presence of mRNA in the vaccine, triggering a type 1 interferon reaction known to favor extrafollicular response(*38, 39*).

### SARS-CoV-2 mRNA vaccination induces higher CSR, but lower somatic hyper mutation in S1^+^ clonotypes

Exploiting the integrable nature of the data in this resource, we matched the single-cell and bulk datasets by identifying sequences with identical CDRH3 amino acid sequences (**Fig. 2A**). We leveraged the sequencing depth in the bulk data (a total of 3,778,590 BCR sequences, collapsed into 1,517,840 unique clonotypes) to generate B cell lineage trees while annotating their antigen specificity using single-cell data. Using this method, we identified 2,453 S1^-^ and 113 S1^+^ clonotypes with matches across both datasets. This approach enabled the identification of a greater number of S1^+^ specific sequences for BCR lineage analysis than would be possible by relying solely on scRNA-seq.

**Figure 2.**
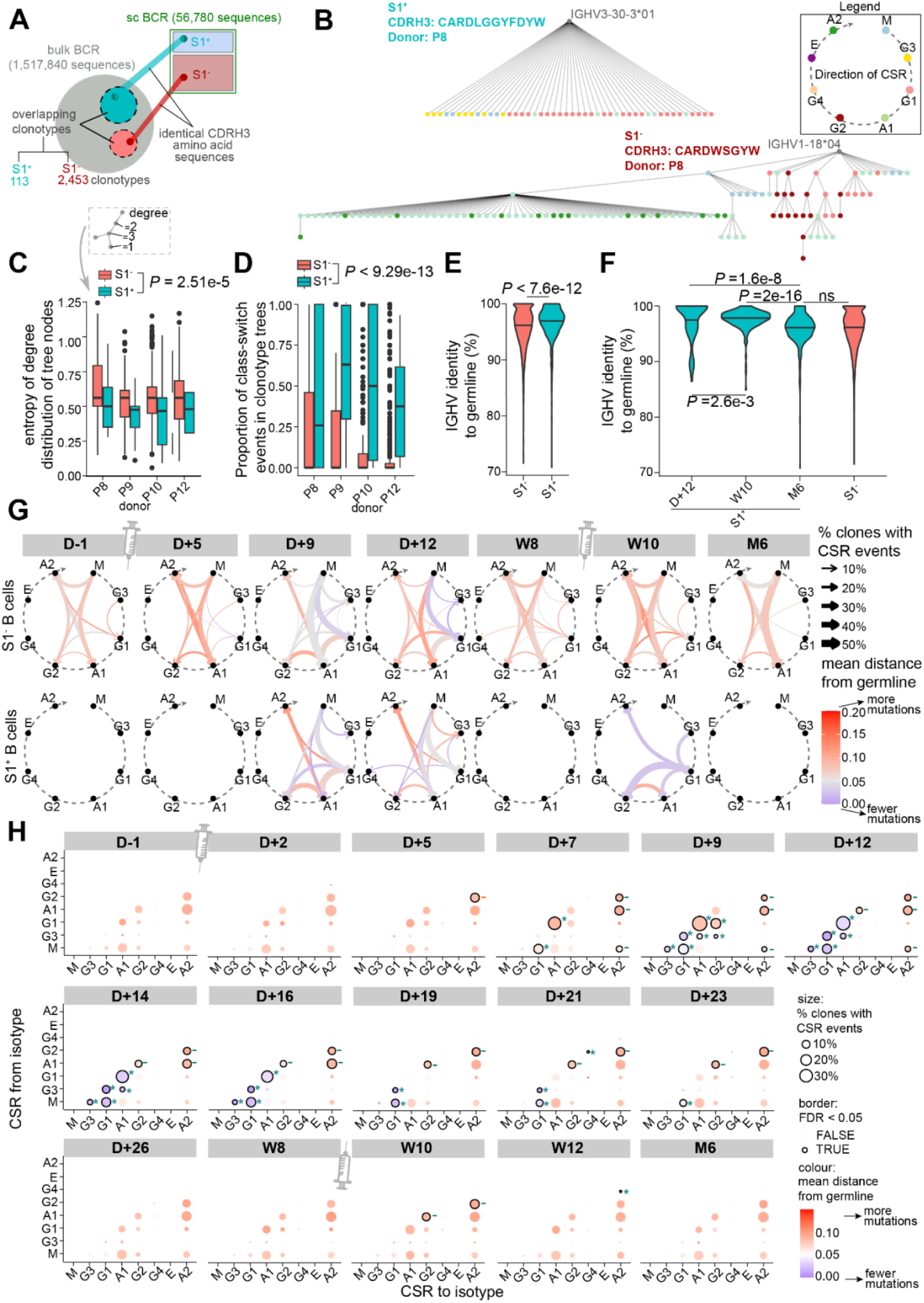
Class-switch recombination occurs in a partially multistep manner with a block at *IGHG2* together with low somatic hypermutation in S1^+^ B cells of naive individuals after SARS-CoV-2 mRNA vaccination. (A) Schematic illustrating matching of single-cell and bulk BCR sequences, based on identical CDRH3 amino acid sequences, to define S^+^ and S^-^ clonotypes sampled in both assays. (B) Representative example trees for S1^+^ and S1^-^ clonotypes identified based on the approach illustrated in (A). Each dot represents an individual BCR sequence, color-coded by their subclasses. Branches connecting nodes of different color denote isotype switching and tree depth denotes acquisition of mutations. (C) Comparison of S1^+^ and S1^-^ clonotype trees in terms of entropy of degree distribution in the trees, as a proxy of the extent of branching events in clonotype trees (Fig. 2B). Only donors with at least 10 S1^+^ clonotype trees were considered in this analysis. Statistical evaluation was performed using a mixed-effect model with antigen specificity (S1^+^/S1^-^) as the fixed effect and donor identifiers as random effects. (D) Frequency of class-switch branches in clonotype trees (Fig. 2B) connecting sequences with different isotypes from S1^+^ and S1^-^ clonotype trees. Only donors with at least 10 S1^+^ clonotype trees were considered in this analysis. Statistical evaluation was performed using a mixed-effect model with antigen specificity (S1^+^/S1^-^) as the fixed effect and donor identifiers as random effects. (E) Comparison of BCR somatic hypermutation between S1^+^ B cells and S1^-^ B cells. Statistical comparisons were based on Wilcoxon’s rank-sum test and p-values were corrected for multiple test corrections based on the Benjamini-Hochberg method. (F) Comparison of BCR somatic hypermutation of S1^+^ B cells between D+12, W8 and M6, and S1^-^ B cells at all timepoints. Statistical comparisons were based on Wilcoxon’s rank-sum test and p-values were corrected for multiple test corrections based on the Benjamini-Hochberg method. (G) Class-switch trajectories in S1^-^ (top) and S1^+^ (bottom) clonotype trees, expressed as a carousel of BCR isotypes arranged clockwise, matching the physical organization of the human *IGH* gene locus. Arrows connect the start and end points of class-switching, with their width proportional to frequency of class-switch events and color depicting the mutational level at which class-switching was estimated to occur. (H) Class-switch events captured in bulk BCR clonotype trees across time points, enumerated separately for the start and end isotypes based on the reconstructed clonotype trees. Size of bubbles correspond to the class-switch frequency involving the specified pairs of isotypes, and colors correspond to the mean sequence distance from the annotated germline at which the class-switch event occurs, as estimated using the reconstructed trees. A higher distance from germline indicates elevated mutational level. Class-switch events with significant (false discovery rate [FDR] < 0.05) frequency changes compared to D-1 are highlighted with black borders. Asterisks (*) denote significant elevated class-switch frequency compared to D-1, and minus signs (-) denote significant decreased class-switch frequency.

We observed distinct structural differences in the lineage trees reconstructed for S1^+^ and S1^−^ clonotypes (**Fig. 2B**). S1^+^ lineage trees exhibited fewer branches and sub-branching events, as quantitatively reflected by reduced entropy in the degree distribution across nodes (i.e., less variation in the number of connections per sequence) (**Fig. 2C**). In addition, and although CSR events were evident in both cases, S1^+^ clonotypes displayed a higher proportion of class-switch event branches in lineage trees compared to S1^−^ clonotypes (**Fig. 2D**). This was accompanied by lower SHM levels in S1^+^ compared to S1^−^ BCR sequences (**Fig. 2E**). SHM levels in S1^+^ B cells did eventually increase at M6 compared to D+12 and W10, becoming comparable to the SHM levels in S1^−^ B cells (**Fig. 2F**). Bulk BCR sequencing further supported this observation, revealing lower mutation rates during the first peak (D+9/D+12), particularly in IgG isotypes (Supp. Fig. 3A).

In summary, despite prior assumptions linking CSR and SHM in human B cell development, our findings indicate that class switch recombination in S1^+^ B clonotypes occurs with minimal evidence of SHM in the first 10 weeks of a primary response.

### Class-switch recombination follows a multistep pattern up to the *IGHG2* gene locus in S1^+^ B cells during SARS-CoV-2 mRNA vaccination

As S1^+^ B cells mainly mapped onto switched B cells (**Fig. 1E-F**) and S1^+^ clonotypes showed higher class-switch branches (**Fig. 2D**), we aimed to explore CSR across different timepoints to reveal its directionality and dynamics.

We started to observe switching in the bulk repertoire as early as day 5, where we could see the relative levels of IgG1 and IgG2 change (Supp. Fig. 3). However, it was not until day 9 that the clone sizes were large enough for us to sample switching in S1^+^ clones. Switching events peaked at D+12 and W10 after the first and second vaccine doses, respectively, with no CSR events observed at M6 (**Fig. 2G**). S1^+^ clones generally followed a stepwise CSR pattern. During the early response (D+9/D+12), S1^+^ clones predominantly switched from *IGHM*, *IGHG3* and *IGHG1*, whereas switching after the second dose (W10) originated mainly from more downstream subtypes such as *IGHG1* and *IGHA1* (**Fig. 2G**). Most switching events occurred towards the two genomically adjacent *IGHC*, such as from *IGHM* to *IGHG3* or *IGHG1*, and from *IGHG1* to *IGHA1* or *IGHG2* (**Fig. 2G**). Occasional switching beyond *IGHG2*, including transitions from *IGHA1* or *IGHG1* to *IGHA2*, was also observed (**Fig. 2G**). As previously noted (**Fig. 2E-F**), these CSR patterns in S1^+^ B cells were accompanied by lower levels of somatic hypermutation compared to S1^−^ throughout the vaccination regimen (**Fig. 2G**). In contrast, S1^-^ clones displayed a more stochastic CSR pattern, which remained largely consistent throughout vaccination. This included the same *IGHG2* blockade, increased switching to IgA2, and higher mutation rates compared to S1^+^ clones (**Fig. 2G**).

Bulk BCR sequencing validated these findings, confirming the *IGHC* expression changes (Supp. Fig. 3B-C), the preference for switching to genomically proximal *IGHC* genes, the rare switching beyond *IGHG2* (**Fig. 2H**), and the lower mutation rates in switching clones (Supp. Fig. 3A).

These observations reveal that antigen-specific clones induced during primary responses in naive vaccinees predominantly follow a "partially multistep" CSR mechanism. In addition, most switch events occur stepwise to subtypes in close genomic proximity, with CSR generally progressing up to *IGHG2*. In view of the importance of IgA2 in mucosal immunity it is notable that the switching route to IgA2 is most often via IgA1 in the early response.

### SARS-CoV-2 mRNA vaccine induces prominent IgG1 but limited IgA class switching

Following the confirmation of a partially multistep CSR pattern (**Fig. 2G**) during SARS-CoV-2 mRNA vaccination, we assessed how this pattern translated into the expression levels and distribution of the different BCR subtypes. Using scRNA-seq and flow cytometry datasets we quantified productive *IGHC* transcripts and surface BCR respectively.

Following the first dose we observed an absolute number and a relative proportional enrichment in *IGHG1*, *IGHA1* and *IGHA2* at D+12, however, in response to the second dose (W10), *IGHG1* followed by *IGHG2* become the dominant subtypes expressed in S1^+^ B cells paralleled by a contraction of *IGHA1* and *IGHA2* (**Fig. 3A-C**). *IGHG1* remained dominant by M6 with reduction of *IGHG2* compared to W10 (**Fig. 3A-C**).

**Figure 3.**
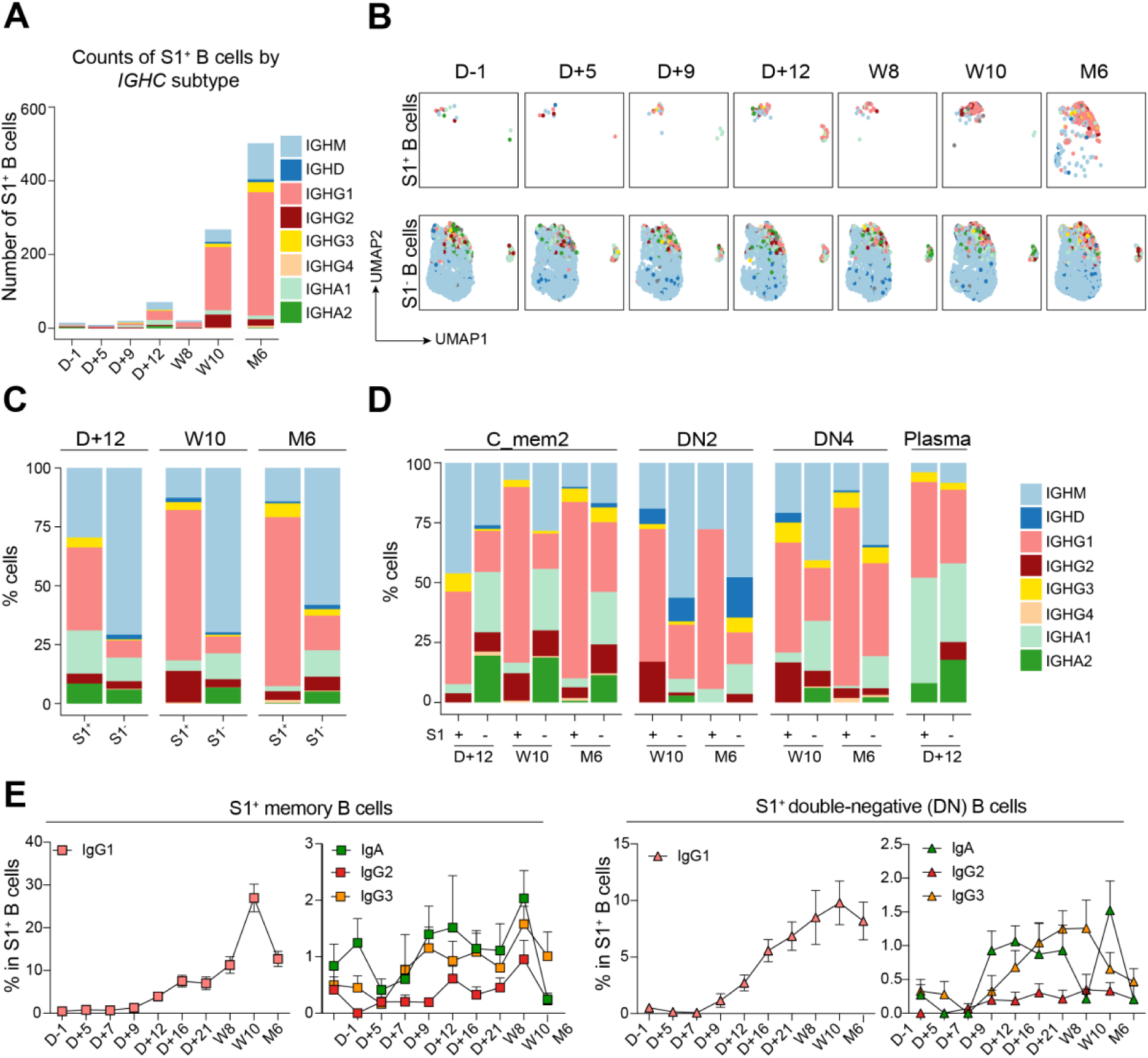
SARS-CoV-2 mRNA vaccine induces prominent IgG1 but limited IgA class switching in S1^+^ B cells. (A) Number of S1^+^ B cells grouped by BCR subtype across time points. (B) BCR subtypes of S1^+^ and S1^-^ B cells color-coded in the UMAP visualization across time points. (C-D) Comparison of BCR subtypes distribution in S1^+^ and S1^-^ B cells at D+12, W10 and M6 time points, over (C) all cells in the scRNA-seq data and (D) separately quantified for C_mem2, DN2, DN4 and plasmablasts subsets. Data is only shown if there are more than 10 S1^+^ B cells for the specified time point/cell subset combination. (E) Frequencies of S1^+^ class-switched memory B cells (CD19^+^CD27^+^CD24^+^CD38l°IgD^-^IgM^-^, squares) and switched double-negative (DN) B cells (CD19^+^CD27^-^IgD^-^IgM^-^, triangles) as percentage of S1^+^ B cells grouped by BCR isotype quantified using flow cytometry data during vaccine response. Data points and trend lines indicate mean over all donors with available data and error bars indicate the standard errors of mean.

Analysis by B cell subpopulations showed that S1^+^ C_mem2, DN2, and DN4 cells expressed higher levels of *IGHG1* but much lower levels of *IGHA1* and *IGHA2* than equivalent S1^-^ B cell subsets at all timepoints (**Fig. 3D**). Notably, the expression of *IGHA1* and *IGHA2* was the highest in S1^+^ plasmablasts compared to the other S1^+^ B cell subsets at D+12 (**Fig. 3D**). Flow cytometry confirmed these findings, showing that IgG1^+^ S1^+^ memory (CD27^+^IgD^-^CD24^+^CD38^lo^) and IgG1^+^ DN (CD27^-^IgD^-^) B cell levels remained predominant, especially at W10, while levels of IgA, IgG2 and IgG3 were limited (**Fig. 3E**, Supp. Fig. 6A-C). Additionally, S1^+^ plasmablasts were the main S1^+^ B cell subset expressing IgA at D+12, confirming the scRNA-seq data (Supp. Fig. 6D). By M6, the frequencies of all switched Ig subtypes within memory B cells, DN B cells, plasmablasts, and total B cells were decreased compared to W10 in S1^+^ B cells, with IgG1 being the main remaining Ig subtype (**Fig. 3E**, Supp. Fig. 6A-E). S1^-^ total, memory, DN B cells and plasmablasts remained largely unchanged during the study (Supp. Fig. 6F-I).

These observations matched with serum antibody titers and BCR bulk repertoire data, showing an expansion of IgG in serum with minimal IgA presence (Supp. Fig. 1B), and an increase in IgG1 compartment in the bulk repertoire following the initial vaccination (Supp. Fig. 3B-C).

These data show that, despite observing multistep switching to isotypes downstream of IgG1, most of the vaccine-derived antigen-specific clones were IgG1. Indeed, high IgG1 expression has been observed by others during COVID-19(*40, 41*) and SARS-CoV-2 vaccination(*42*) as well as during other viral infections such as Ebola, influenza, and RSV(*8, 43–47*). Furthermore, mRNA SARS-CoV-2 vaccination induced poor IgA responses which were mainly detectable in plasmablasts and only during the first peak of the response.

### *IGHG2* blockade is bypassed by S1^+^ DN and C_mem2 B cells expressing specific productive *IGHC* transcripts

After observing a reduction in multistep switching beyond the *IGHG2* gene by both scBCR-seq (**Fig. 2G**) and bulk BCR-seq (**Fig. 2H**), we sought to further investigate this phenomenon by comparing the levels of productive and sterile *IGHC* transcription in S1^+^ and S1^-^ B cells (**Fig. 4A**). In accordance with the elevated switching observed in S1^+^ clonotypes (**Fig. 2D**), general sterile transcription (x-axis), a proxy for CSR directionality(*28*), was higher in S1^+^ B cells compared to S1^−^ B cells (**Fig. 4A**). Similarly, and in consonance with the class-switching events (**Fig. 2G-H**), both productive and sterile *IGHC* transcripts displayed the previously observed partially multistep CSR pattern with a block in sterile transcription beyond the *IGHG2* gene in both S1^+^ and S1^−^ B cells (**Fig. 4A**). Exceptions to this arrest included B cells expressing productive *IGHG4* for S1^−^ B cells (**Fig. 4A**). As control for S1^−^ B cells, analysis of a previously published scRNA-seq data of non-vaccinated healthy control B cells(*12*) shows a similar pathway, confirming that the arrest at *IGHG2* is independent from vaccination (**Fig. 4B**).

**Figure 4:**
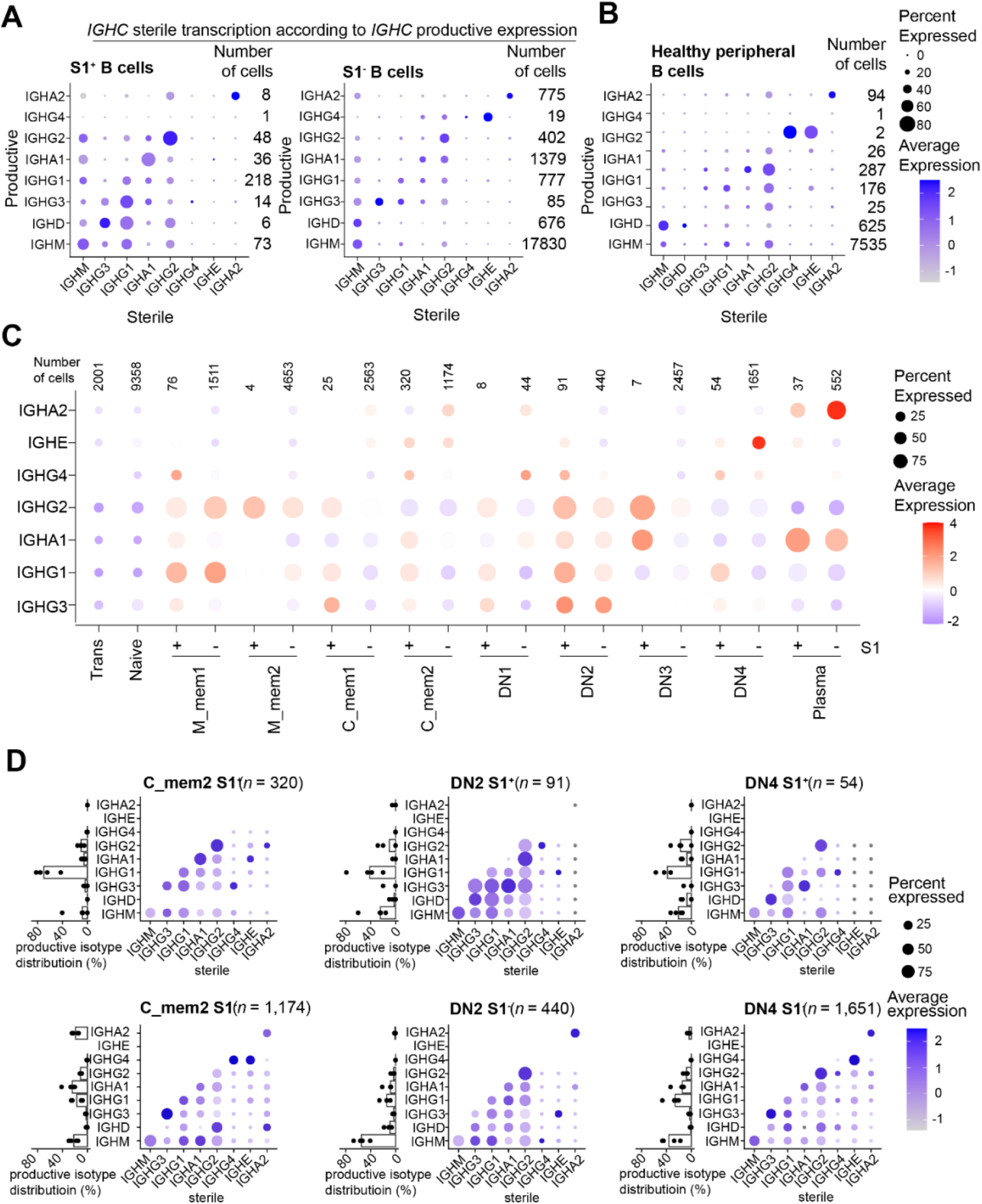
*IGHG2* blockade is bypassed by S1^+^ DN and C_mem2 B cells expressing specific productive *IGHC* transcripts. (A-B) Quantification of S1^+^ and S1^-^ B cells in (A) our single-cell (sc) transcriptomic data and (B) a reference dataset of peripheral B cells at homeostasis from Stewart et al.(*12*), in terms of their productive isotype (vertical axis, based on sc BCR-seq data) and sterile transcript expression (horizontal axis, quantified using sciCSR). Dot sizes are proportional to amounts of cells positive for the given sterile transcripts and color depicts expression level. The numbers of B cells expressing each productive transcript are indicated. (C) Quantification of sterile transcription level separately for S1^+^ and S1^-^ cells sampled up to W10 in each B cell subset. (D) Quantification separately for S1^+^ and S1^-^ cells in the C_mem2, DN2 and DN4 subsets, in terms of their productive BCR isotype distribution (left, bar plot) determined using sc BCR-seq data, and sterile transcription levels for B cells of different BCR isotypes (right, dot plot). For bar plots, data points correspond to individual donors.

Next, we hypothesized that distinct B cell subsets might exhibit different sterile transcription patterns and hence only certain subsets might be able to bypass the *IGHG2* blockade. We identified DN subsets, C_mem2, and plasmablasts as the primary subtypes expressing sterile transcripts beyond the *IGHG2* locus (**Fig. 4C**).

Furthermore, as observed above (**Fig. 4A**), we asked whether sterile transcription was associated with the productive transcription of *IGHC* at a B cell subset specific level. Indeed, S1^+^ DN2 and DN4 B cells, expressing productive *IGHG1*, *IGHA1* and *IGHG2*, but not *IGHG3*, were able to progress beyond the *IGHG2* locus barrier expressing sterile *IGHG4* and *IGHE,* with minimal *IGHA2* sterile expression (**Fig. 4D**). Similarly, S1^+^ C-mem2 B cells expressing productive *IGHG3*, *IGHA1* and *IGHG2* but not *IGHG1* co-expressed *IGHG4*, *IGHE* and *IGHA2* sterile transcripts respectively (**Fig. 4D**). The distribution for S1^−^ B cells was more complex and promiscuous with a wider range of *IGHC* subtypes expressing sterile transcripts beyond *IGHG2*. Particularly interesting was the *IGHE* sterile expression by S1^-^ DN4 and C_mem2 B cells expressing several different productive transcripts, especially *IGHG4* (**Fig. 4D**).

This is the first evidence that sterile transcription beyond the *IGHG2* locus is dependent on a combination of B cell subset and BCR subtype.

### Bulk BCR sequencing reveals a more diverse B cell response but fails to highlight many antigen-specific cells

A successful B cell response requires both class switch recombination and the optimization of antigen specificity governed by variable (V), diversity (D) and joining (J) segment usage in the BCR repertoire complemented by SHM. Repertoire studies make the logical assumption that identifying expanded clones in a bulk BCR repertoire is a proxy for identifying antigen-specific cells. The data resource here includes 3.8 million sequences of high-quality long reads across all timepoints, enabling a broad analysis of VDJ gene usage and expansion.

Relative to the baseline (D-1), we observed a transient increased representation of several V, D and J genes, e.g. *IGHV3-33*, *IGHV3-30*, *IGHV*3-13, *IGHD1-26* and *IGHJ6* during the first peak (D+9/+12) suggesting a polyclonal primary response (**Fig. 5A**). However, only the *IGHV4-34* gene, previously found to be overexpressed in hospitalized COVID-19 patients(*8*), was upregulated in both primary and secondary response peaks (W10) (**Fig. 5A**). Of interest, we observed that changes in VDJ gene usage varied between BCR subtypes (Supp. Fig 7A).

**Figure 5:**
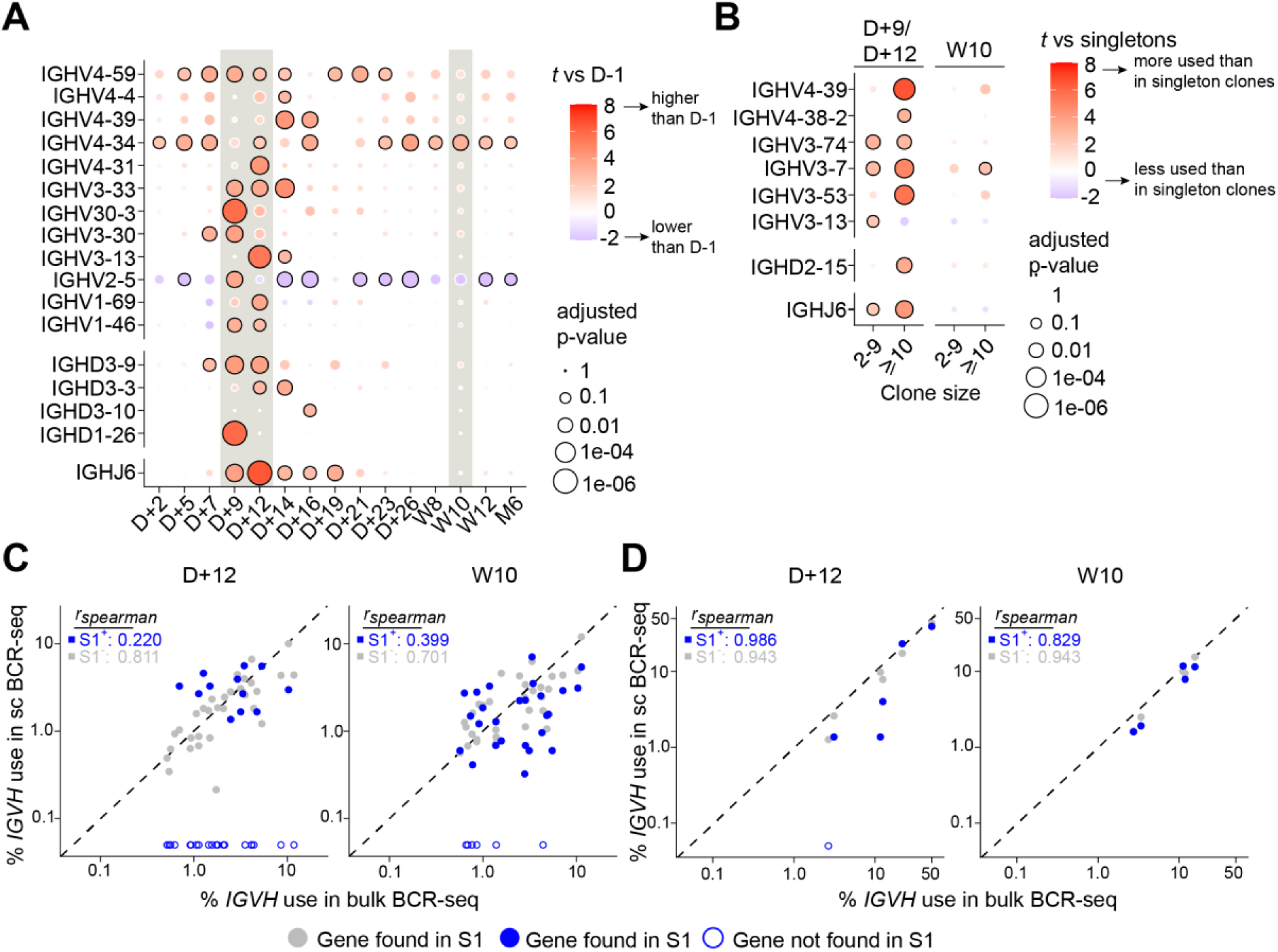
Differential polyclonality in VDJ gene usage during first versus second peak and direct comparison of single-cell and bulk BCR sequencing data. (A) Change in immunoglobulin heavy-chain variable (V), diversity (D) and J (joining) gene usage across timepoints in the bulk BCR sequencing data. Gene usage was evaluated on subsets of BCR sequences which are mutated (<99% sequence identity to the germline). First (D+9 and D+12) and second (W10) response peaks are boxed in grey. Statistical significance was assessed by fitting mixed-effect linear models of percentage gene usage (dependent variable) against time point as the fixed effect and donor identifiers as the random effect. Bubble color depict effect size compared to D-1 (positive values in red indicate elevated usage of gene compared to D-1), and bubble sizes correspond to p-value after false-discovery rate adjustment. (B) Comparison of heavy-chain V, D and J gene usage between clonotypes of different sizes. Gene usage was computed for sequence subsets as defined in (A), separately for time points at the first peak (D+9 and D+12) and the second peak (W10). Clonotypes were grouped by their sizes, into singletons, clonotypes with 2-9 sequences and those with more than 10 sequences (“>=10”). Mixed effect models were fitted to compare gene usage against singleton clonotypes as a control, with clonotype sizes as fixed effects and donors as random effects. Bubble color depict effect size (positive values in red indicate elevated usage of gene), and bubble sizes correspond to p-value after false-discovery rate adjustment. (C-D) Comparison of heavy-chain (C) V gene and (D) J gene usage estimated using the bulk BCR sequencing [BCR-seq] (X axis) and single-cell (sc) BCR-seq (Y axis). Each data point represents gene usage for an individual *IGHV* or *IGHJ* gene in a single donor. For single-cell BCR data, gene usage was quantified separately for S^+^ and S^-^ B cells. Genes not detected in single cells are represented as empty circles. Spearman correlation (*r_spearman_*) between the gene usage in bulk BCR-seq data and S1^+^ and S1^-^ B cells were indicated.

Comparison of heavy-chain V, D, and J gene usage across expanded clonotypes of different sizes (2-9 versus >10 sequences per clonotype) revealed the preferential usage of certain genes according to clone size. Among others, the *IGHV3-53* gene, frequently associated with SARS-CoV-2 neutralizing antibodies(*48*), was notably enriched in larger clonotypes (**Fig. 5B**). As before, this phenomenon was also associated with BCR subtypes (Supp. Fig 7B).

Additionally, we found that the Gini index, a measure of diversity, of larger clonotypes, increased over time, peaking at M6. Diversity of smaller clonotypes (2-9 sequences) remained unaltered during the vaccination regime (Supp. Fig. 7C).

Complementarity Determining Region 3 (CDRH3) of *IGH* genes encodes a crucial antigen-binding region of the antibody and can often be seen to be altered at the population level in response to immune challenge. Bulk BCR sequencing data showed increased length and hydrophobicity in the CDRH3 regions of *IGHG1* sequences at D+12 (Supp. Fig. 8A-B), which aligned with previous findings(*8*), and correlated with RBD-specific IgG titers (Supp. Fig. 8C).

The integrable nature of our data enables comparison between bulk repertoire data and data on antigen positive cells from scBCR-seq. The CDRH3 analyses are in agreement: BCR *IGHV* sequences of S1^+^ had significantly longer CDRH3 regions than S1^−^ B cells (Supp. Fig. 8D). When compared by *IGHC* subtype, S1^+^ expressing *IGHG1* and *IGHG2* exhibited longer CDRH3 regions than their S1^−^ B cells (Supp. Fig. 8E). Similarly, CDRH3 regions in S1^+^ B cells that express the *IGHG1, IGHG2, IGHM* and *IGHA1* subtypes are more hydrophobic compared to the same CDRH3 regions in S1^−^ B cells (Supp. Fig. 8F).

There are, however, some interesting points of divergence between the single-cell and the bulk-repertoire BCR datasets which highlight the importance of considering sampling in the interpretation of such studies. Of the 1,297 S1^+^ sequences in the enriched scBCR-seq data, only 113 (8.7%) were found in the bulk data. The background matching, when cells were not enriched for antigen specificity, was 2,453 out of 34,335 (7%) S1^-^ cells matched with sequences in the bulk repertoire. Similarly, there are expanded clonotypes in the bulk repertoire that are not evident in the scBCR-seq. Mapping the frequency of *IGH* germline gene use between the two types of data shows a good correlation for S1^-^ scBCR-seq data for both *IGHV* and *IGHJ* gene usage (**Fig. 5C,D**), but the correlation statistic decreases for S1^+^ scBCR-seq *IGHV* gene usage. It should also be noted that in addition to sampling considerations, the 3.8 million sequences in the bulk repertoire may reflect a skewed view of “expanded clonotypes” since plasma cells express, on average, 100-fold more immunoglobulin transcripts than B cells in our single-cell data (Supp. Fig. 8G).

## Discussion

Our study provides a unique multi-immunomic resource for studying the dynamics of the human primary immune response. The intensive sampling schedule, particularly within the first three weeks of the initial vaccine dose, has enabled unparalleled temporal resolution of early B cell dynamics, BCR repertoire, and CSR. This approach provides new insights into the evolution of the immune response, offering a deeper understanding of B cell development compared to previous studies(*49–58*). We have used these data to finely map the temporal evolution of a robust B cell response, integrating the data from serology with that from flow cytometry, bulk BCR repertoire, scRNA-seq and scBCR-seq to compare critical events and peaks of responses between observational methods and revealing paradigm-shifting insights into the regulation of antibody class switch recombination (CSR).

SHM and CSR have been tightly linked as both require the activation induced cytidine deaminase (AID) enzyme, thought to be restricted to the germinal center(*59*). However, there is a disconnect between these two processes as we observe CSR long before accumulation of typical levels of SHM. New evidence in mice shows that CSR occurs at the follicle edge and is decoupled from SHM which would support our observations(*28*). Classically, from studies in mice, it has been thought that primary T cell-dependent immune responses showed early accumulation of hypermutation (around 7 days post-immunization)(*60, 61*). In contrast, we found that hypomutation persisted throughout most of our sampling time course in S1^+^ B cells, including at the peak of the secondary response (W10). Hypomutation of switched S1^+^ B cells has also been previously reported in COVID-19 disease(*62*). One possible explanation is because S1^+^ B cells do not need SHM. However, this is not likely the case since by month 6 the SHM levels are comparable to S1^-^ B cells. Recent observation of fine needle aspirate samples from human lymph nodes also shows a slow and prolonged affinity maturation in the human germinal center(*63*).

Antibody class switching without SHM could suggest an extrafollicular response by S1^+^ B cells in the early stages as B cells that have not entered the germinal center usually show low mutation rates(*64–66*). The nature of the vaccine, containing mRNA, could be relevant here since RNA can trigger a type 1 interferon reaction known to favor extrafollicular responses(*38*). The early expansion of DN2 B cells would concur with this possibility(*37*). Indeed, several studies report extrafollicular responses and increased DN during COVID-19 and SARS-CoV-2 vaccination(*67–70*). Antigen-specific DN2 were greatly downregulated in the long term, while mutation rates increased in S1^+^ B cells, which could be attributed to the development of canonical follicular and GC responses. The matching of the month 6 S1^+^ cells with Ig genes in bulk data only after day 12 (data not shown) would also fit with there being a two-stage response.

Organization of the BCR data into lineage trees enables the study of CSR events as they occur along the timeline. Integrating the scBCR-seq with bulk data overcomes the limitations that occur with the data types individually, allowing exploration of specificity as well as the volume of data needed to provide sufficient lineages. The steady state picture at D-1 is similar to that published by Horns et al(*71*), acknowledging that switching can occur both directly from IgM to another subclass, or can switch between subclasses. However, a static picture misses intermediate stages and these new data enable a distinction between background steady state and antigen labeled (by scRNA-seq) or likely antigen specific (as inferred by appearance of hypomutated sequences) over a detailed time series. The new picture that emerges is of a “partial multistep” switching, with some direct routes and some routes that proceed via closer intermediates in a 5’ to 3’ direction as the immune response proceeds. Furthermore, in primary responses from naive individuals the switching began from upstream isotypes such as IgM, IgG3 and IgG1, but secondary responses had downstream originators such as IgG1 and IgA1. Switching beyond IgG2 along the locus is limited to IgA2, and it is very rare to see IgM to IgA2 transitions, rather IgA2 would mostly arise via an IgA1 intermediate. This information is crucial for researchers who aim to elicit IgA2 responses, for mucosal vaccination for example.

The paradigm-shifting discovery in these data is the openness of the *IGHC* locus to enable multistep switching, but only up to the *IGHG2* gene. This was not just apparent in the productive transcripts of the lineage trees, but also in the patterns of sterile transcription in the scRNA-seq data. Thus, it appears that activation of B cells causes sterile transcription of all *IGHC* genes as far as *IGHG2*, and only in certain circumstances will transcripts occur past this point. These circumstances likely involve multiple developmental changes in a B cell, since we find that the exceptions to the *IGHG2* blockade (in both S1^+^ and S1^-^ B cells) were in particular types of B cell: the DN and C-mem2 subsets. Moreover, when subdividing these cell types further, by the expression of productive Ig transcripts, we found only certain B cell subsets expressing specific BCR isotypes expressed sterile transcripts beyond *IGHG2*. This is the first study to observe these phenomena in such detail and these results are key for understanding the immune response and CSR dynamics in naive individuals.

The diversity of the BCR response to the vaccine after the first dose is considerable. The separate sampling events that occurred to provide the bulk BCR repertoire and the scBCR-seq repertoire have highlighted that there are potentially many other antigen-specific cells that are missed. Only 17% of our S1^+^ single cell clones matched with the clonotypes from bulk sequencing, and many of the clonally expanded sequences in the bulk data were not seen in the S1^+^ single cells. The incongruence of observations at the bulk and single-cell levels has implications in using bulk BCR repertoire as a tool for the characterization of antibody responses.

In summary, we provide a unique multi-omic resource that can serve as a blueprint to analyze CSR and other aspects of B cell response during primary human immune challenges. Our observations revolutionize the way we think about the dynamics of CSR, showing generalized activation of CSR up to *IGHG2* with B cell development and Ig-subtype-specific states allow breaching of the block at *IGHG2*. This resource could be used to build a generalizable model of CSR by comparative analysis against other vaccination and infection studies in naive individuals in the future. Our scRNA-seq data serves as an atlas of B cell states for discovering new genes involved in the regulation of CSR and other aspects of B cell maturation, and as a general resource to interrogate proven CSR regulating molecules and pathways identified in animal models. Identifying similarities and differences in clonal dynamics, analyzing how CSR processes intersect with BCR clonal diversification and affinity maturation, together with integrating these findings with experimental or computational assessments of antigen specificity in different settings will significantly advance our knowledge of human B cell activation. This is required to improve vaccines that can cause switching beyond IgG to favor IgA1 and IgA2 for greater mucosal protection(*72–74*), or to be applied to better understand and optimize CSR in a disease-specific manner such as for autoimmunity and cancer immunotherapy.

A limitation of our methodology is the sampling bias, particularly in single-cell transcriptomics, where we cannot sample all B cells and clones. This means that we may have missed intermediate steps of some of the direct switching events we observed, overestimating the proportion of direct switches. Sampling bias is an important aspect to note in all studies involving BCR repertoire sampling with our data highlighting the limited overlap one can get in matching samples from the same person, even in the larger S1^-^ scBCR-seq data there was a low overlap of clonotypes between the single cell and bulk datasets. In addition, we sampled peripheral immune cells without assessing tissue residency. Consequently, the lower presence of IgA^+^ S1^+^ B cells could be due to a migration of these to mucosal surfaces. However, the evidence does not support this as studies have found that SARS-CoV-2 infection, but not intramuscular vaccination, elicits IgA^+^ B cell residency(*75*). Indeed, we corroborated this as we did not observe IgA secretion in saliva of vaccinees but in COVID-19 patients. Finally, some features of the observed immune response may be attributable exclusively to the mRNA vaccine platform such as the increase of DN B cells and BCR hypomutation, as other studies, using protein-(*76, 77*), vector-based(*78–83*), or inactivated vaccine platforms(*84, 85*) have not investigated or found the presence of DN or atypical memory B cells in humans.

## Materials and Methods

### Study design and participants

This study was designed to investigate the immune response to the SARS-CoV-2 vaccine with particular focus on CSR and B cell responses in a highly systematic timeline to analyze immune processes in a dynamic way. Fifteen healthy adults between 24 and 35 years of age were immunized with the mRNA-1273 vaccine formulating mRNA molecules encoding for the full length of the spike (S) protein of the original SARS-CoV-2 strain. The participants were immunized during the spring of 2021 as part of the national (United Kingdom) vaccination program. Demographic details of the participants can be found in Supplementary Table 1. Informed consent was asked of all participants prior to the start of the study. Participants were surveyed for previously known SARS-CoV-2 infection and other relevant (immune-related) co-morbidities at the start and at the end of the study. Ethical approval was obtained from the East Midlands - Leicester Central Research Ethics Committee, under REC reference no. 21/EM/0064. Additionally, COVID-19 infection samples were collected from SARS-CoV-2 positive patients at Frimley and Wexham Park hospitals during 2020 (consented under UK London REC no. 14/LO/1221)(*8*).

Whole blood, serum, peripheral blood mononuclear cells (PBMCs), saliva and nasal wash were taken for a variety of readouts including measurement of antigen-specific antibody levels by ELISA, antibody blocking capacity, multicolor spectral flow cytometry for immunophenotyping from whole blood and PBMCs, single-cell RNA sequencing (scRNA-seq) and bulk B cell receptor (BCR) sequencing at various time points (**Fig. 1A**). A baseline time point (Day −1) was scheduled 24h prior to initial vaccination which happened on day 0 with subsequent timepoints happening every Monday, Wednesday and Friday for 3 weeks: time points D+2 to D+26 (**Fig. 1A**). The second vaccine dose visit occurred 8 weeks after the first dose with a baseline time point (week 8 or W8) taken before this second immunization. Additional two post-second dose timepoints at 10 (W10) and 12 (W12) weeks after initial vaccination, or 2 and 4 weeks after second dose respectively, were taken, with a final time point 6 months (M6) after initial vaccination (**Fig. 1A**). Participants were screened for previous SARS-CoV-2 reactivity (either through infection or cross-reactivity) with two participants (P6 and P14) showing anti-RBD IgG antibodies at baseline (Supp. Fig. 1A). These two participants were excluded from subsequent analysis.

For single-cell transcriptomics, the following time points were selected for analysis: D-1 as baseline, D+5 which was prior to class-switched B cell expansion (Supp. Fig. 3B-C), as well as D+9 and D+12 which coincided with changes in IgG and IgA class-switched antibodies at the secreted and transcriptional levels (Supp. Fig. 1B and Supp. Fig. 3B-C). Cells from W8 and W10 were also analyzed to characterize secondary B cell responses to the vaccine.

### Whole blood, PBMC and serum isolation

Whole blood was collected in sodium heparin tubes (455051, Greiner Bio-One) and serum SST tubes (456018, Greiner Bio-One). 100ml of non-coagulated whole blood from heparin tubes was kept separately for whole blood *ex vivo* immune phenotyping. Non-coagulated whole blood from heparin tubes was diluted 1:1 in PBS (14040-091, Gibco) with 2% heat-inactivated fetal bovine serum (FBS, FCS-SA/500-22512, Labtech). Density gradient centrifugation using SepMate tubes (85450, STEMCELL Technologies) was used to isolate peripheral blood mononuclear cells (PBMCs) according to the manufacturer instructions. PBMCs were counted and viability estimated using a trypan blue exclusion assay. Cells were then resuspended at 10×10^6^ viable cells per ml in FBS containing 10% DMSO (20688, ThermoScientific) and stored in liquid nitrogen. Serum SST tubes were centrifuged at 1200g for 10 minutes at room temperature, then serum was aliquoted into cryovials and stored at −70 °C.

### Quantification of RBD-specific antibody titers

Anti-RBD IgM/A/G ELISAs were performed based on a published plasma-based method(*86*). High-binding 96 well plates (9018, Corning for COVID Vaccine samples and 442404, ThermoFisher for COVID patient samples due to plastic shortages) were coated with 100µl of 2μg/ml of RBD (ab273065, Abcam) in PBS and incubated at 4°C overnight. Plates were then washed three times with 0.01% Tween 80 (P5188-100ML, SigmaAldrich) in PBS (PBS-T) and blocked with 200μl of 3% milk (84615.0500, VWR) in PBS-T for between 1 and 4 hours. Blocking buffer was removed and plates tapped dry, and 100ml of diluted samples/controls added. Serum was diluted at either 1:20, 1:60, 1:100, 1:300, 1:900 or 1:2700 with PBS to have the OD fall within the linear range of the standard curve/positive controls and diluted once more 1:3 on the plate with PBS-T with 1% milk powder. Positive controls for corning plates were loaded at initial concentrations of 0.2 ng/μl for IgM (Ab01680-15-0, Absolute Biotech, clone CR3022), 0.602 ng/ml for IgA (Ab01680-16-0, Absolute Biotech, clone CR3022), 0.2 ng/μl for IgG (ab273073, Abcam, clone CR3022) and for maxisorb at 0.06 ng/ml for IgM, 0.20067 ng/ml for IgA, 0.06 ng/μl for IgG, and then serially diluted 1:2 four times. All samples and positive controls/standard curve points were run in duplicate. Each plate also contained two blanks and negative controls from pre-pandemic serum samples. Samples were incubated for 2 hours at room temperature, washed 3 times and 50 ml of HRP-conjugated detection antibody (IgM: A18835, ThermoFisher; IgA: A0295-1ML, Sigma-Aldrich/Merk; IgG: A18817, ThermoFisher) added and incubated for 1 hour (RT), then washed 3 times and loaded with 100 ml of OPD (#11879250, FisherScientific) to reveal for 15 minutes and finally with 50 ml of 3M Hydrochloric acid (HCl) used to stop the reaction. Plates were read on a SpectraMax iD3 plate reader (Molecular Devices) at 490 nm. Sample values were standardized against the blank control by subtraction and antibody titers interpolated in GraphPad Prism (v9) and multiplied by their dilution factor.

### Bulk B cell receptor (BCR) library generation

3ml of whole blood at each timepoint was taken into Tempus Tubes (4342792, Applied Biosciences) and RNA was extracted according to manufacturer’s instructions. Bulk Immunoglobulin repertoire libraries were prepared as previously described(*8*). Briefly, a 5’ template switch transcription was performed to incorporate Unique Molecular Identifiers (UMIs), followed by two rounds of polymerase chain reaction (PCR). The first PCR stepped-out to add a primer landing site at the 5’ end, facilitating the step-out addition of donor identifier barcodes at the 5’ end in PCR2 for multiplexing. The reverse primers were designed to nest in the constant regions with step-out multiplex identifiers added in PCR2. Libraries were sequenced on a Pacific Biosciences (PacBio) Sequel IIe system at the Liverpool Centre for Genomic Research. Quality control, data cleaning and removal of multiplicated UMIs were performed as previously described(*8*). Our bulk BCR repertoire dataset contained a total of 3,778,590 sequences.

### Blocking assay

A V-PLEX SARS-CoV-2 panel 30 (ACE2) kit (K15635U, Meso Scale Diagnostics) was used according to the manufacturer’s instructions to measure the capacity of serum samples to block the binding of variants of SARS-CoV-2 to the ACE2 receptor. A dilution of 1:50 was used for all serum samples.

### Whole blood *ex vivo* immune phenotyping

100ml of non-coagulated fresh whole blood from heparin tubes was stained with additional 100µl staining buffer the surface antibodies described in supplemental methods table 1 at room temperature in the dark for 30 minutes. Then 2ml of red blood cell lysis and fixation buffer (00-5333-54, eBioscience) were added and incubated at room temperature in the dark for 25 minutes. Fixed samples were centrifuged at 800g for 5 at room temperature. Finally, cells were washed with 3ml of FACS buffer (PBS 2% FBS and 0.2mM EDTA (15575-038, Invitrogen)) and centrifugation at 800g for 5min at room temperature. Samples were acquired in a Digital LSR II flow cytometer (BD Biosciences). Flow cytometric data were analyzed using FlowJo software (v10.8.1). Antibodies for whole blood staining were previously specially titrated for this staining. A higher concentration than that used for PBMCs was chosen as antibody concentration used for PBMCs yielded poor staining. This is likely due to the presence of red blood cells in great numbers in whole blood samples.

**Supplemental methods table 1:**
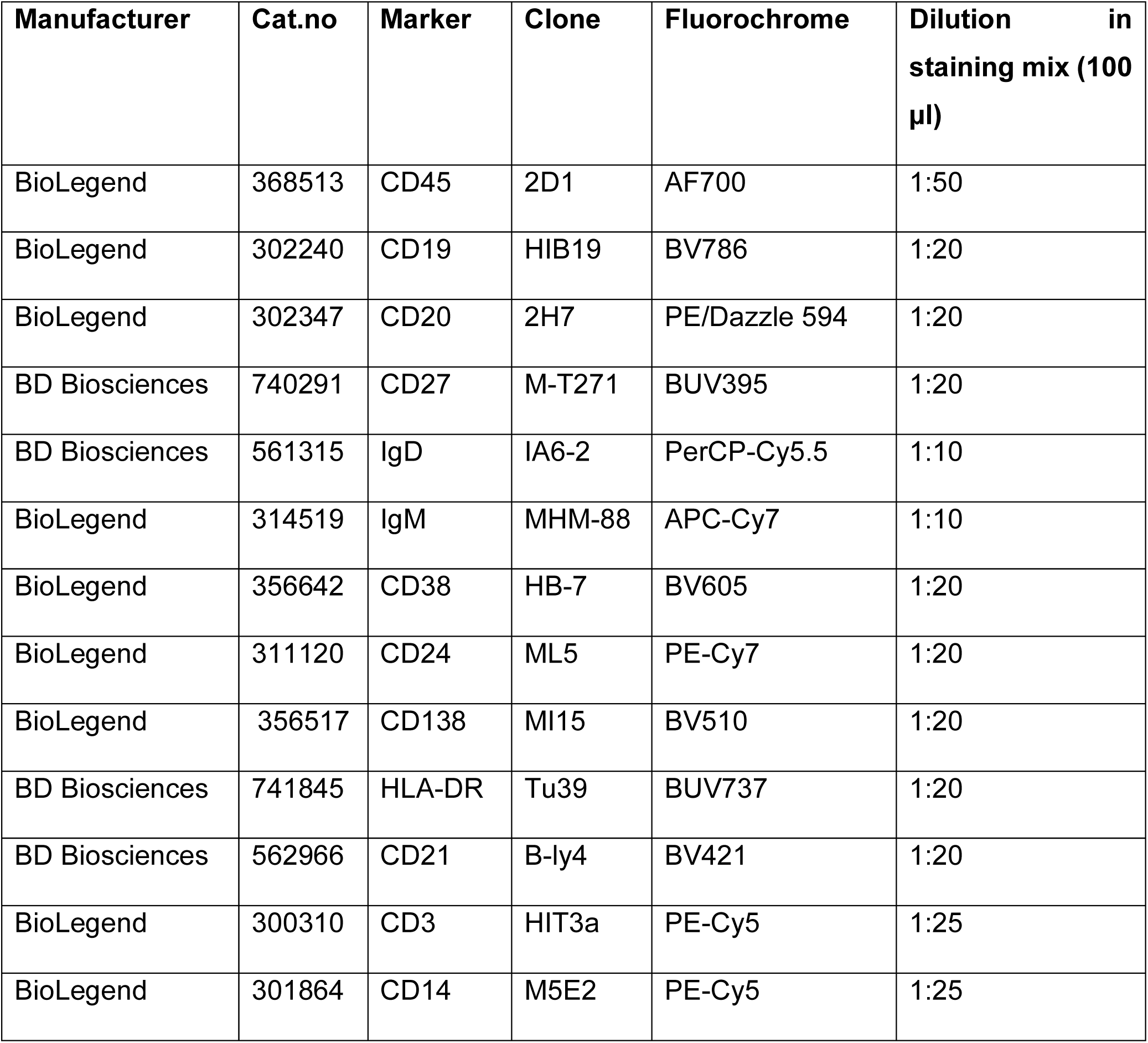
whole blood *ex vivo* immune phenotyping surface antibodies.

### Antigen-specific B cell *ex vivo* phenotyping

Vaccine-derived antigen-specific B cells were identified by tagging them with the subunit 1 (S1) of the spike protein (S) of the ancestral SARS-CoV-2 and the receptor-binding domain (RBD), a domain contained within the S1. Biotin-conjugated S1 was coupled with two fluorochrome-labelled streptavidin to form two different S1-fluorochome conjugates. Biotin-S1 (793806, Biolegend) was conjugated with streptavidin-BV421 (405225, Biolegend) and streptavidin-APC (405207, Biolegend) separately, in PBS at a ratio of 1:6 (streptavidin-fluorochrome:biotin-S1). Biotin-conjugated RBD (793904, Biolegend) was conjugated with streptavidin-BUV737 (612775, BD Biosciences) in PBS at ratio of 1:4 (streptavidin-fluorochrome:biotin-RBD). A decoy conjugate consisting of only biotin coupled with a streptavidin-fluorochrome complex was constructed by mixing D-biotin (B-4501, SigmaAldrich) and streptavidin-PE-Cy5 (405205, Biolegend) at a ratio of 1:40 (streptavidin-fluorochrome:free D-biotin). These conjugates were incubated under agitation at 4°C for at least one hour. 5 x 10^6^ PBMCs were stained per sample (for each participant and timepoint analyzed). PBMCs were thawed at 37°C until a small amount of ice remained, 500μl of pre-warmed 37°C FBS was added, samples transferred to a 15ml falcon and diluted to 10ml with complete RPMI (10% heat-inactivated FBS and 1% penicillin/streptomycin (P0781, SigmaAldrich), centrifuged at 500g for 8 minutes at 4°C and washed once with PBS by centrifuging at 500g for 8 minutes at 4°C. Cells were incubated in 200µl of LIVEDEADÔ fixable blue (L23105, Invitrogen) diluted 400-fold in PBS together with FcR blocking agent (130-059-901, Miltenyi) at 4°C in the dark for 30 minutes. Cells were then washed with FACS buffer and centrifuged at 500g for 8 minutes. 200µl of 5mM D-biotin FACS buffer (D-biotin FACS buffer) containing 5ng of Biotin-PE-Cy5 (decoy) was added and incubated for 30 minutes in the dark at 4°C. Following incubation with the decoy, cells were washed twice with D-biotin FACS buffer by centrifuging at 500g for 8 minutes. Cells were then stained with the antigen probe cocktail (0.5mg/samples for each S1 construct and 0.25mg/sample for the RBD construct) and surface antibodies (described in supplemental methods table 2) in 200ml of D-biotin FACS buffer and left in the dark at 4°C for one hour to incubate. Tubes containing the cells were agitated every 20 minutes during this incubation period to ensure maximal staining. Cells were then washed with D-biotin FACS buffer and centrifuged at 500g for 8 minutes before 200µl of IC Fixation Buffer (00-8222-49, eBioscience) were added left to incubate for 20 minutes at 4°C in the dark. Following fixation, cells were washed twice with FACS buffer by centrifuging at 500g for 8 minutes and resuspended in FACS buffer for acquisition. Samples were acquired in a CytekTM Aurora cytometer (5 lasers) using SpectroFlo (v3.0.3) with automated unmixing. Flow cytometric data were analyzed using FlowJo software (v10.8.1). In total we analyzed 16,293,828 S1^-^ and S1^+^ B cells over all donors and timepoints considered in this flow cytometry analysis.

**Supplemental methods table 2:**
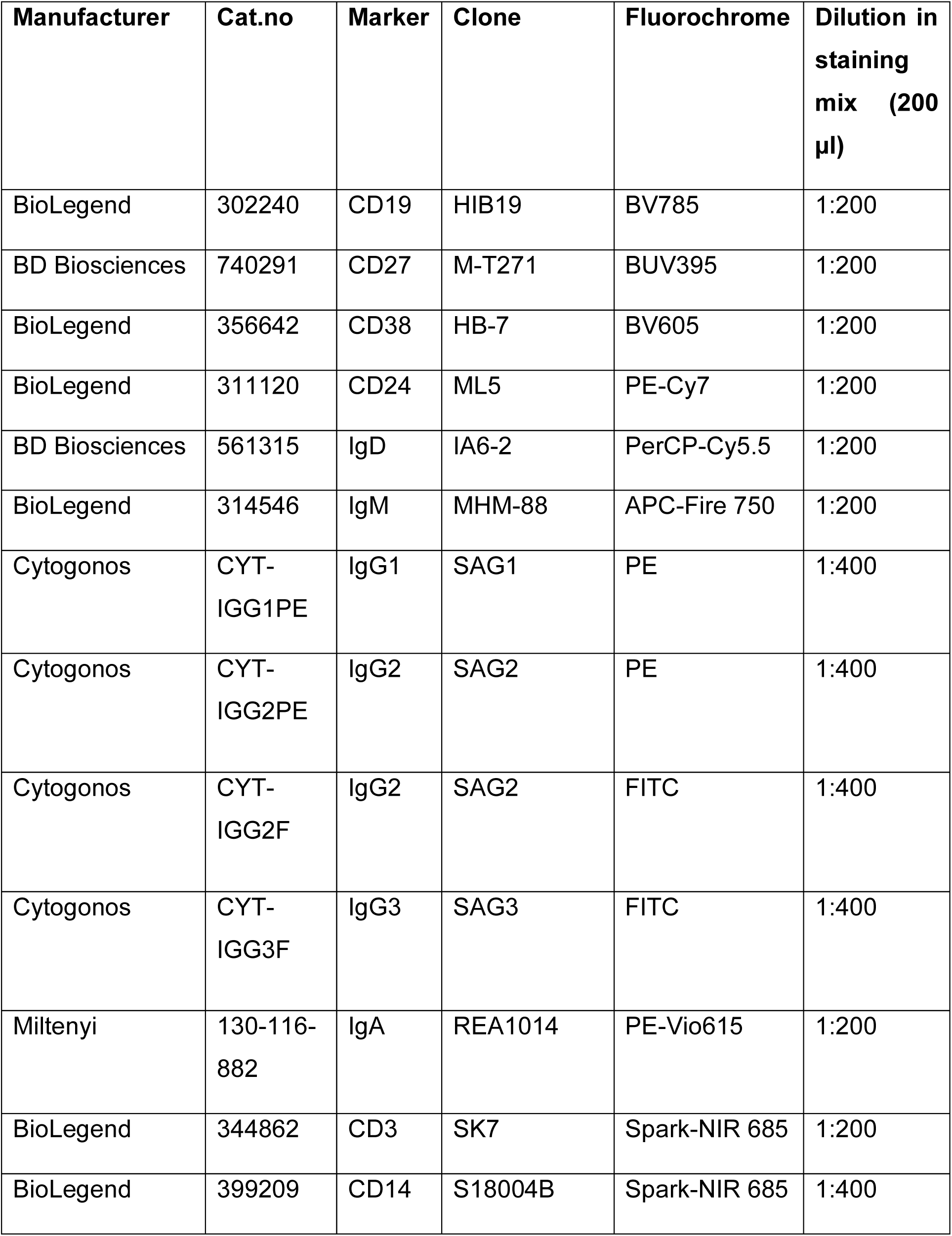
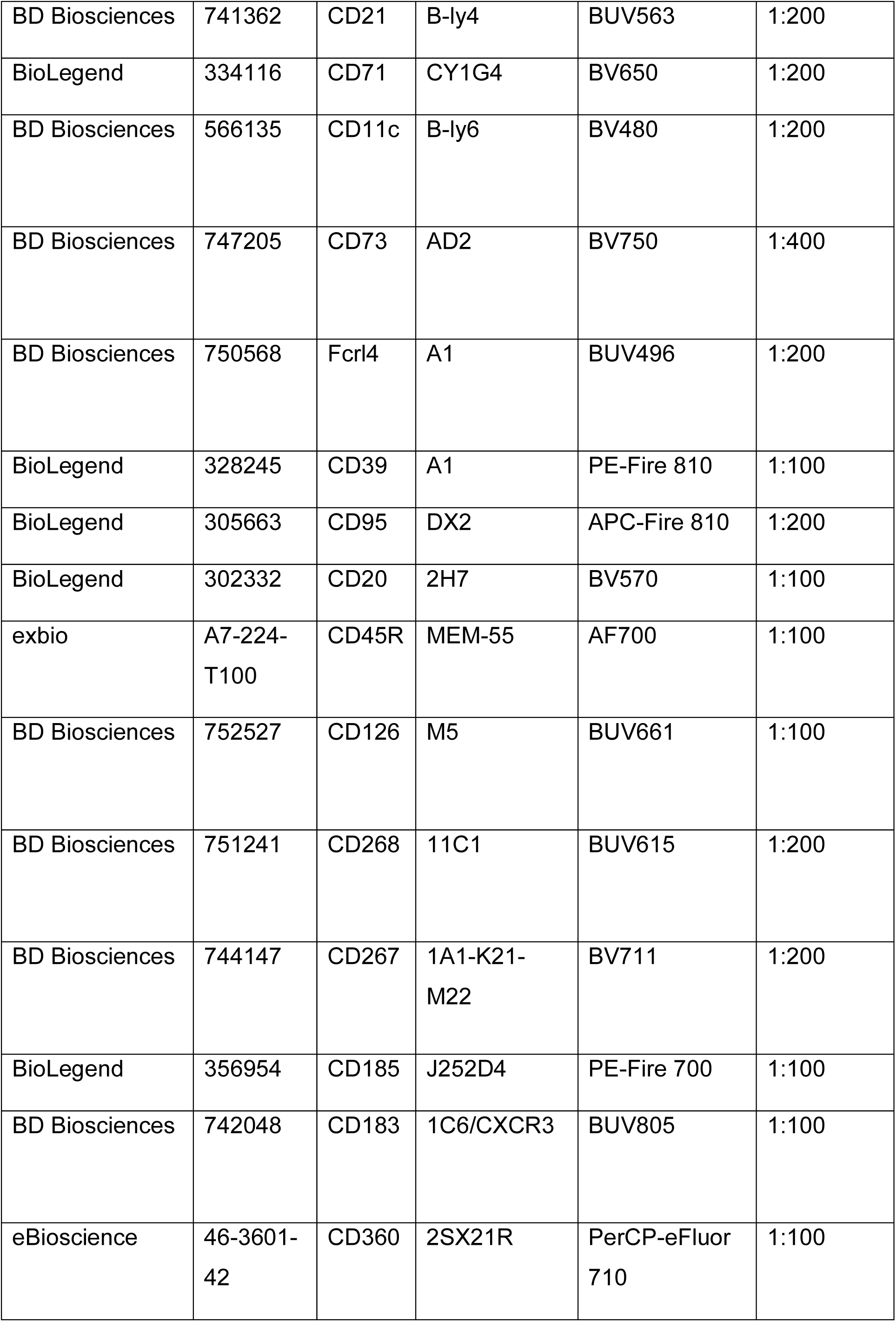
*Ex vivo* B cell phenotyping surface antibodies for antigen-specific B cell *ex vivo* phenotyping.

### Single-cell transcriptomic library generation

Vaccine-derived antigen-specific B cells were identified by tagging them with the subunit 1 (S1) of the spike protein (S) of the ancestral SARS-CoV-2 and the RBD. Biotin-conjugated S1 was coupled with two fluorochrome/oligomer dual-labelled streptavidin to form two different S1-fluorochome/oligomer conjugates. In this way, antigen-specific B cells were sorted via flow cytometry assisted sorting (FACS) using the fluorochromes and sequenced using single cell technologies using the oligomer sequence with a posterior bioinformatic identification of antigen-specific B cells. Biotin-S1 (793806, Biolegend) was conjugated with Totalseq™ streptavidin-PE (405261, Biolegend) and Totalseq™ streptavidin-APC (405283, Biolegend) separately, in PBS at a ratio of 1:6 (streptavidin-flurochrome+oligomer:biotin-S1). Biotin-conjugated RBD (793904, Biolegend) was conjugated with streptavidin-oligomer (405271, Biolegend) in PBS at ratio of 1:4. A decoy conjugate consisting of only biotin coupled with a streptavidin-fluorochrome complex was constructed by mixing D-biotin and streptavidin-FITC (405201, Biolegend) at a ratio of 1:40 (streptavidin-FITC:free D-biotin). These conjugates were incubated under agitation at 4°C for at least one hour. 5 x 10^6^ PBMCs were stained per sample (for each participant and timepoint analyzed). PBMCs were thawed at 37°C until a small amount of ice remained, 500μl of pre-warmed 37°C FCS was added, samples transferred to a 15ml falcon and diluted to 10ml with cRPMI and counted, washed once with PBS by centrifuging at 500g for 8 minutes at 4°C. Cells were incubated in 200µl of staining mix containing Zombie NIR (423105, Biolegend), both S1 construct (S1-APC+oligomer, S1-PE+oligomer) the RBD construct (RBD-FITC), the FITC decoy (D-biotin-FITC) and the extracellular antibodies described in supplemental methods table 3 at 4°C in the dark for 1 hour.

**Supplemental methods table 3:** Flow cytometry assisted sorting surface antibodies.

| Manufacturer | Cat.no | Marker | Clone | Fluorochrome | Dilution in staining mix (200 µl) |
| --- | --- | --- | --- | --- | --- |
| BioLegend | 302240 | CD19 | HIB19 | BV421 | 1:200 |
| BioLegend | 367141 | CD14 | 63D3 | BV786 | 1:200 |

Cells were then washed with FACS buffer and centrifuged at 500g for 8 minutes at 4°C. Samples were acquired and sorted in a BD FACSAria™ Fusion. All events were gated for lymphocytes based on FSC/SSC, singlets and living cells. B cells were identified as CD19^+^ cells, decoy negative B cells further selected, and S1^+^ B cells detected by double PE and APC staining (Supp. Fig. 2D). Three populations were sorted: S1^+^ B cells (CD19^+^PE-S1^+^APC-S1^+^), S1^-^ B cells (CD19^+^PE-S1^+^APC-S1^+^) and CD19^-^ lymphocytes. Innate cells were gated on the same basic criteria and size selected based on size with CD14 on z-axis to aid size selection. Samples across time points for each donor were processed in the same batch and sorted cell populations from each time point were pooled in equal numbers, with hashtags added individually for each day’s sample (Biolegend 294661, 394663, 394665, 394667, 394669, 394671) to allow bioinformatic demultiplexing using the feature barcode sequencing reads. A pool of all S1^+^ B cells, 60% S1^-^ B cells, 20% CD19^-^ Lymphocytes, 20% Innate cells was run in 4 10X reactions lanes for 5000 cells in each reaction. Cells were centrifuged (500g 5 minutes at 4°C) and resuspended in PBS with non-acetylated BSA to the desired concentration and run on the 10X chromium controller utilizing the Chromium Next GEM Single cell 5’ reagent kit v2 (Dual Index) (document number: CG00030 Rev F) producing the GEX, VDJ sequencing and cell surface protein libraries according to the manufacturer’s instructions. Libraries were sequenced on a HiSeq2000 or NovaSeq X Plus Series (PE150) at 50,000 reads per cell for GEX libraries and 5,000 read per cell for VDJ/Cell surface protein libraries.

### BCR repertoire data analysis

BCR sequences were annotated for immunoglobulin VDJ gene usage using IMGT/HighV-Quest(*87*). Clonotype clustering was performed as previously described(*8*) by calculating Levenshtein distance pairwise between CDRH3 nucleotide sequences. The resultant distance matrix was hierarchically clustered, and branches were cut at 0.05 to define clones. Physicochemical properties were calculated using the R Peptides package (*88*). Clonal diversity was calculated using the Gini coefficient, which measured the evenness in the distribution of clone size. We used the transformation (1 – Gini coefficient) as a measure of clonal diversity.

BCR lineage trees were constructed using the BrepPhylo package(*8*). Briefly, a maximum parsimony tree was first constructed for each clone using the dnapars executable in the phylip package(*89*), using the IMGT-gapped V-gene nucleotide sequences as input. All clones with at least 3 sequences were considered. From these trees we calculated, for each observed sequence in a given clone, its distance to the annotated germline gene. This tree-based distance from the germline measures the extent of mutation accumulation for the given sequence (*8*). We further analyzed the reconstructed lineage trees to identify class-switch events, i.e., branches in the tree which connect BCR sequences of different isotypes. In BrepPhylo we previously implemented routines to prune the dnapars trees (which were built using only V gene sequences) to remove edges which implicate CSR events that violate the physical order of constant region genes in the human IGH locus, and build minimum spanning arborescence tree of the pruned data using Edmond’s algorithm (*8*). These trees were used to identify and quantify CSR events between any pairs of isotypes in the data.

### Single-cell transcriptomic data analysis

#### Data preprocessing and clustering

Matching 10X genomics gene expression, BCR and feature barcode/cell surface protein libraries were processed through CellRanger multi version 6.1.2. The following reference genome versions were downloaded from the cellranger website for sequence alignment and annotation: refdata-gex-GRCh38-2020-A (for gene expression libraries) and refdata-cellranger-vdj-GRCh38-alts-ensembl-5.0.0 (for BCR libraries). For data processing steps outlined below, the R package Seurat (v4.3.0)(*90*) was used unless otherwise stated. The raw read count matrix for each library was first preprocessed to collapse individual genes belonging to each of the following groups of genes to eliminate donor-specific variations from dominating downstream cell clustering: immunoglobulin variable (V) genes (gene name patterns matching regular expression “^IG[HKL]V[0-9]”), diversity (D) genes (“^IG[HKL]D[0-9]”), joining (J) genes (“^IG[HKL]J[0-9]”), ribosomal genes (“^RP[LS]|^MRP[LS]”), individual HLA class Ia genes (“^HLA-[ABC]$”), HLA class Ib genes (“^HLA-[EFG]$”), HLA class II genes (“^ HLA-D”). Read counts mapped to genes belonging to these gene groups were collapsed into separate metagenes and replaced the individual genes listed therein. The percentage of reads mapped to mitochondrial genes (“^MT-|^MTRNR”) per-cell were calculated and appended as cell metadata. Cells with transcripts mapped to between 200 and 4000 distinct genes & a mitochondrial read percentage below 15% were retained for analysis. The SCTransform(*91*) protocol implemented in Seurat was applied for read count normalization, with the mitochondrial read percentage modelled as a covariate to remove variation attributable to this factor. Genes at the immunoglobulin, T cell receptor, HLA and mitochondrial loci were removed from the list of variably expressed genes prior to dimensionality reduction and clustering to remove their impact in driving the definition of cell clusters. Dimensionality reduction was performed using principal component analysis (PCA) using this pruned list of variably expressed genes. A k-nearest neighbor graph was constructed using the first 14 principal components, and cell clustering was performed on this graph using the FindClusters function with the resolution parameter of 0.5. B cells were identified by examining CD20 (*MS4A1*) expression across these clusters. The B cell clusters were subsetted from the data and analyzed separately from the non-B cells. For B cells, cell labels were assigned per cell by transferring from our previously published scRNA-seq atlas of peripheral B cells (*12*), using the TransferData protocol implemented in Seurat, by projecting the PCA structure of the reference to this dataset to classify cells based on similarity in this (projected) PCA space.

#### Demultiplexing

Time point specific hashtags were demultiplexed following the HTODemux protocol implemented in Seurat, where hashtag read counts were normalized using centered log-ratio transformation and thresholds for demultiplexing were determined using the 95^th^ percentile of the normalized read-count distribution as the cut-off to define hashtag positive and negative cells.

#### Identifying S1-specific B cells

Initial analysis of the raw read count data for feature barcodes corresponding to the S1 or RBD baits indicated that signals for the three antigen-specific Totalseq™ feature barcodes (PE Strep-S1, APC Strep-S1, and Strep-RBD) vary greatly, with a lack of signals corresponding to PE Strep-Spike in comparison with Totalseq™ APC Strep-Spike. Totalseq™ Strep-RBD raw counts suffered from significant background noise. We reasoned that an explicit model of background and true signals in these read counts *was* necessary for confident identification of S1-specific B cells. We therefore trained a totalVI(*92*) model using our paired gene expression and feature barcode libraries. TotalVI explicitly model background and foreground distributions for feature barcode libraries(*92*); here we used this to correct for substantial noise in our raw read counts corresponding to the antigen-specific barcodes. We used the get_protein_foreground_probability() function to obtain the corrected antigen-specific signal as a probability score between 0 and 1; the probability distributions for Totalseq™ APC Strep-Spike and Strep-RBD were bimodal with two peaks close to 0 and 1. We therefore used these probability scores to identify S1- and RBD-specific B cells, using a cut-off at 0.5 for identifying antigen-positive B cells.

#### Subclustering of B cell subsets

The three B cell subsets enriched in S1^+^ B cells (C-mem2, DN2 and DN4) were subclustered to investigate gene expression signatures specific to those cells within each B cell subset that were S1-specific. For each B cell subset considered, cells belonging to the specified subset were extracted and subject to variably expressed gene identification, PCA, clustering and UMAP projection using the identical procedure described above for processing the B cell dataset at large, except that the number of principal components considered for UMAP and clustering procedure was chosen separately for each B cell subset such that each included principal component contributed at least 2% of the overall variance. The FindClusters function was used to isolate subcluster(s) specific for S1^+^ B cells within C-mem2, DN2 and DN4 separately, using the resolution parameter of 0.5.

#### Differential gene expression analysis

Differential gene expression was calculated at the pseudobulk level using the scran::pseudoBulkDGE function. This procedure was applied in two contexts: (1) comparison of all S1^+^ versus all S1^-^ cells at Week 10; (2) comparison of S1^+^ versus S1^-^ cells at Week 10 for a given B cell subset. For (1), Week 10 cells with the same combination of the following metadata were aggregated to generate pseudobulk samples: donor identifier, B cell subset label and S1^+^/S1^-^ binary label. For (2), the following metadata were considered to generate pseudobulks: donor identifier, timepoint, subcluster labels. Pseudobulks representing fewer than 3 cells were removed from the comparison. The following formulas were used to construct the design matrix used in the differential gene expression analysis: for (1), ∼ donor + B cell subset + S1 (the coefficient and p-value specific to the S1 term were extracted to generate volcano plots); for (2), ∼ donor + subcluster label (the coefficient and p-value specific to the subcluster enriched in S1+ cells were extracted for volcano plots).

#### Sterile and productive immunoglobulin transcript analysis

We used sciCSR(*14*) to analyze BAM alignments of the gene expression libraries, for identifying sterile transcripts at the immunoglobulin heavy-chain gene locus. Sterile transcripts are immunoglobulin transcripts which lack coding information for the V, D and J gene segments, and instead start at genomic positions 5’ to the beginning of every constant (C) region genes. Sterile transcription is an indication of B cells poised for class-switching(*14*). Using sciCSR we obtained a read count matrix for all heavy-chain sterile transcripts, which were subsequently log-normalized, and the expression levels of these transcripts were visualized using dot plots. For productive transcripts, we integrated the matching single-cell BCR libraries with the gene expression data based on matching cell barcode. In cases where multiple heavy and/or light chain transcripts share the same cell barcode, these cells were flagged and only the transcript with the highest unique molecule identifier (UMI) count was retained for merging with the cell metadata. Columns in the filtered contig annotation output from cellranger corresponding to V, D, J, C genes were retained in the merged cell metadata for BCR isotype-based analysis presented here. V region germline identity was obtained via IgBLAST analysis against the set of germline immunoglobulin VDJ alleles obtained from IMGT (accessed 28-Jul-2022). Physicochemical properties of CDRH3 amino acid sequences were calculated using the Peptides package(*88*) through BRepertoire(*92*).

### Bulk and single-cell BCR data integration

We identified exact overlaps in terms of CDRH3 amino acid sequence between the bulk and single-cell BCR annotated sequence data, and utilize the antigen specificity labelling information in the single-cell data to annotate clonotype lineages sampled in the bulk data. This approach capitalizes on the deep sampling of the antibody repertoire in the bulk libraries whilst filling in antigen specificity information missing from this data. For single-cell data, we considered only sequences with cell barcodes where exactly one heavy and one light chain productive transcript were observed. The CDRH3 amino acid sequences of clonotypes from the bulk data were scanned for exact matches with the single cell data, to extract a list of donor and clonotype identifiers (hereafter “overlapping clonotypes”). This procedure considered only heavy-chain sequences from both the bulk and single-cell data. An overlapping clonotype was annotated as S1^+^ if at least one of the heavy-chain sequences in this clonotype that originated from the single-cell data corresponds to a B cell with S1-binding probability score (see subsection “Identifying S1-specific B cells” under “Single-cell transcriptomic data analysis”) greater than 0.5. CSR-aware arborescence trees were reconstructed for every overlapping clonotype using BRepPhylo (*8*) using the identical procedure as detailed above for bulk BCR repertoire analysis. For each edge in the tree we annotated whether the edge connects sequences of different isotypes to identify class-switch events.

To systematically compare the topologies and class-switch features of the S1^+^ and S1^-^ trees, we enumerated the proportion of branches with class-switch events for each clonotype tree. We also sought to derive a metric to quantify the extent of branching in the trees, to distinguish cases where specific sequences have a large number of direct descendants in the tree versus trees with more sub-branching events (which indicated the acquisition of more mutations at different branches of the tree). To quantify this, we converted each tree into a directed network and enumerated the number of outgoing edges per sequence in the clonotype, i.e., the “out-degree” of each node. We used the entropy of this out-degree distribution as a proxy of branching. These metrics were calculated for each tree and compared between S1^+^ and S1^-^ clonotypes.

### Statistics and data visualization

Statistical analysis was performed in the R statistical computing environment (v4.2.0). Data visualization was produced using the ggplot2 package (version 3.4.1). Mixed effect models were fitted using the lmerTest R package (version) and model estimates were optimized according to the restricted maximum likelihood (REML) criterion. Wherever possible, mixed effect models were fitted with donor as random effect and the fixed effect was set to be the variable for which the desired contrasts were analyzed.

## Acknowledgments

The authors thank the participants of the study for their time and willingness to donate samples during the thorough schedule. The authors thank the members of the Research Facility at the Institute of Immunity and Transplantation (University College London). This work was funded by the Biotechnology and Biological Sciences Research Council (BB/T002212/1 with F.F. as principal investigator). E.S. was funded by a personal PhD fellowship by the University of Surrey. The funders had no role in the collection and analysis of the samples, in the interpretation of data, in writing the report, nor in the decision to submit the paper for publication.

## Author contributions

G.M.G., A. T. S., P. B., D. Kateregga, E.S., C. J. M. P. and Z. B. performed sample collection and processing.

A. G., D. K. D.-W. and C. M. planned sample collection.

G.M.G., J. C. F. N., A. T. S., P. B., F. F., D. K. D.-W., and C. M. designed experiments G.M.G., A. T. S. and E. S. performed experiments.

G.M.G., J. C. F. N., A. T. S., E. S., D. K. D.-W., D. Kipling, D. G. and L. S. analyzed data.

P.B., F. F., D. K. D.-W. and C. M. provided supervision for experimental design and data analysis.

G.M.G., J. C. F. N., A. T. S., F. F., D. K. D.-W. and C. M. prepared the manuscript. All authors read, commented, and approved the final manuscript.

## Resource Availability

Further information and requests for resources and reagents should be directed to and will be fulfilled by Deborah K. Dunn-Walters upon request.

### Materials availability

This study did not generate new unique reagents.

### Data and code availability

All raw data will be deposited at ArrayExpress and will be made publicly available as of the date of publication. Any additional information required to reanalyze the data reported in this paper is available from the lead contact upon request.

## Supplementary materials

**Supplementary Figure 1.**
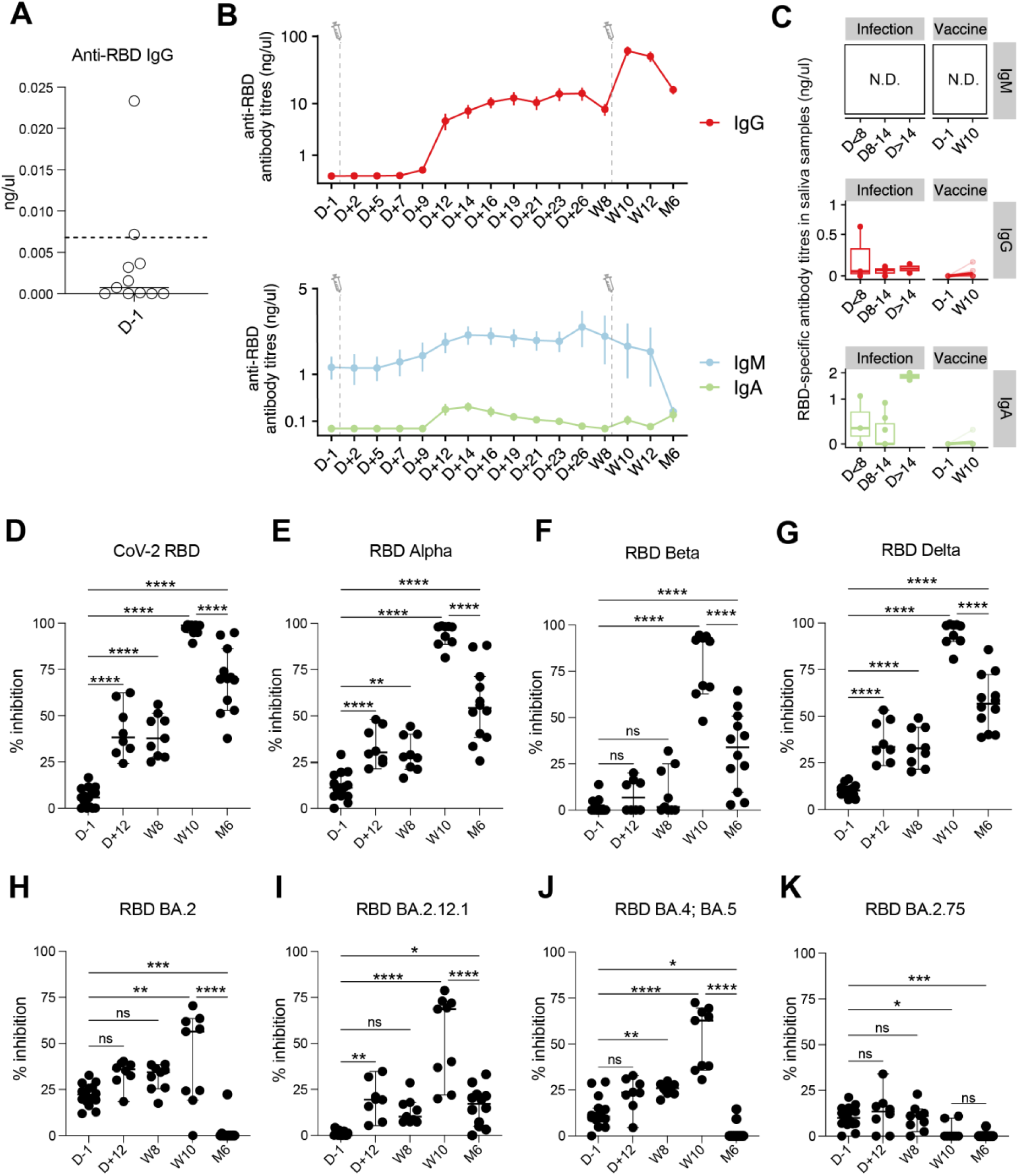
SARS-CoV-2 vaccination against ancestral strain induces antibody titer changes and impairs long term blocking capacity of RBD-omicron. (A) Anti-Receptor Binding Domain (RBD) IgG antibody titers measured at D-1 using ELISA. Data points from two outlier donors with previous SARS-CoV-2 exposure (P6 and P14) were highlighted separately. Dotted line indicate ELISA detection limit. (B) Changes in serum RBD-specific IgG, IgA, and IgM levels across time. Trend line represents mean values per time point across donors with available data; error-bars depict standard error of means. (C) Antibody titers of IgM, IgG, and IgA in saliva samples from hospitalized SARS-CoV-2 infected patients (Stewart, Sinclair, Ng et al. Front Immunol 2021) (“Infection”) and this vaccination cohort (“Vaccine”). For the Infection cohort, time points denote number of days since hospitalization. Data points denote measurements from individual donors. (D-K) Percentage inhibition of binding to the ACE2 receptor against different SARS-CoV-2 strains (D) CoV-2 (Wu-1/ancestral), (E) Alpha, (F) Beta, (G) Delta, and omicron stains (H) BA.2, (I) BA.2.12.1, (J) BA.4;BA.5, and (K) BA.2.75 of serum collected at different timepoints during the vaccine response.

**Supplementary Figure 2.**
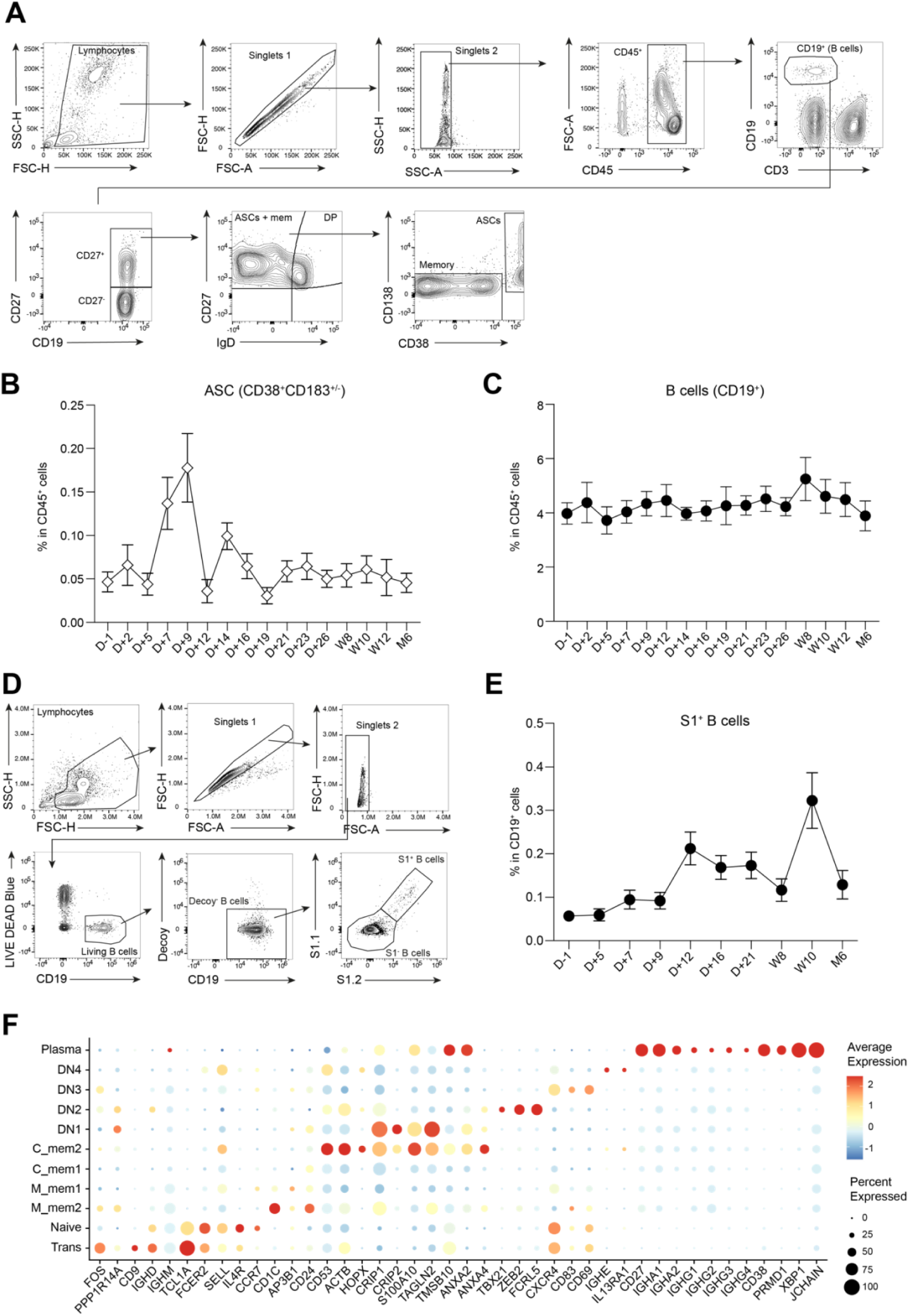
The B cell compartment and S1^+^ B cells in flow cytometry and single-cell BCR sequencing data. (A) Representative plots showing the gating strategy for identification of total CD19^+^ B cells (CD19^+^), and circulating antibody secreting cells (ASCs, CD19^+^CD27^+^IgD-CD38^+^CD138^+/-^) in whole blood. (B) Changes in CD38^+^ antibody secreting cells (ASCs) as percentage of CD45^+^ cells across time. (C) Frequency of total B cells (CD19^+^) as a proportion of CD45^+^ cells during vaccine response. (D) Representative plots showing the gating strategy for identification of antigen-specific (S1^+^) B cells through flow cytometry analysis or flow cytometry assisted sorting (FACS) and subsequent single cell RNA sequencing. (E) Frequency of S1^+^ total B cells (CD19^+^) as a proportion of CD19^+^ cells during vaccine response. (F) Dotplot displaying key marker genes for 11 B cell subpopulations annotated in the scRNA-seq data.

**Supplementary Figure 3.**
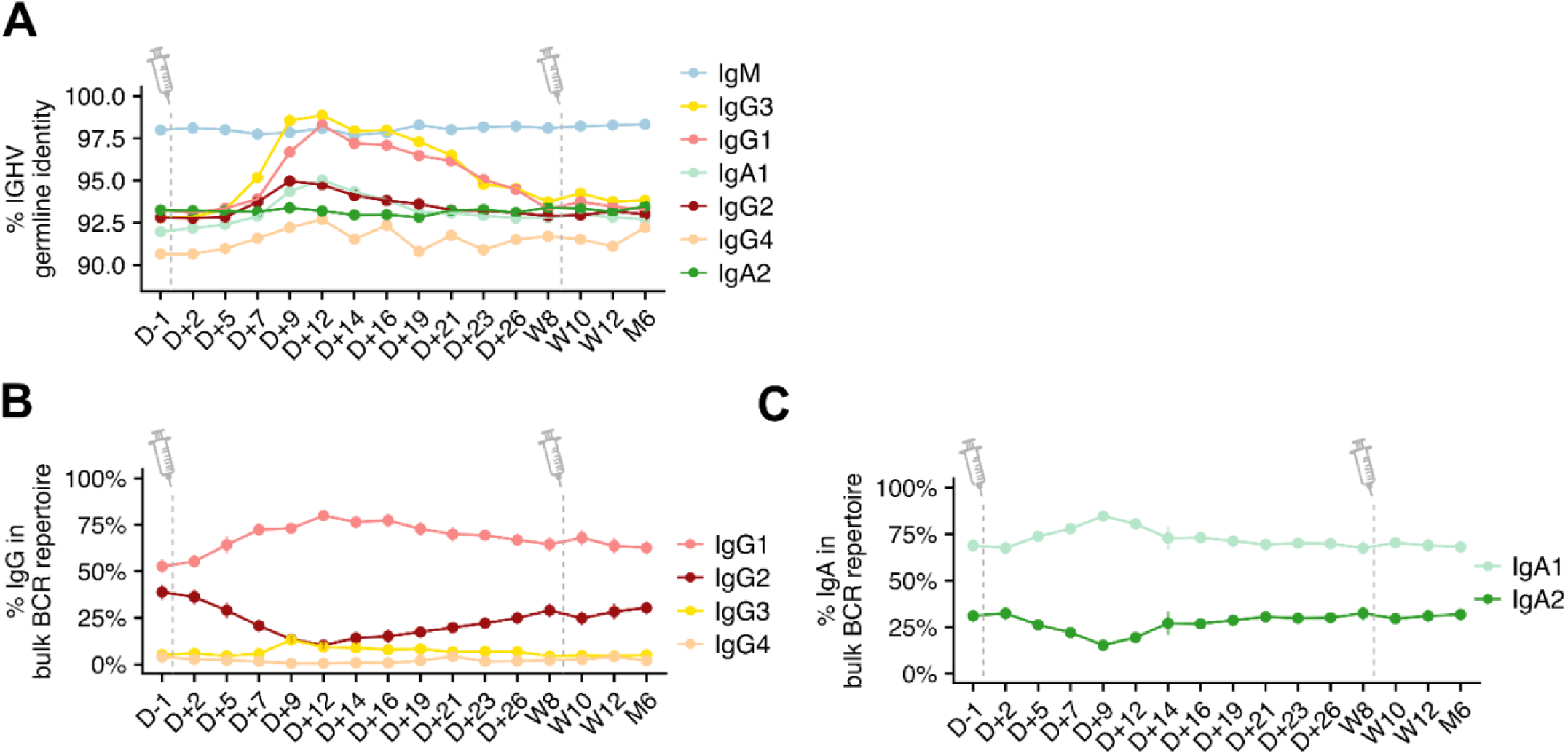
Evolution of *IGHV* germline identity and *IGHC* subtype-proportion in bulk BCR sequencing data. (A) Changes in BCR sequence identity to germline *IGHV* gene divided by isotype subclass across the vaccination time-course. Trend-lines display mean values per time point across all donors with available data; error-bars depict standard error of means. (B-C) Changes in subclass percentage distribution based on bulk B cell receptor (BCR) repertoire data for (B) IgG isotypes and (C) IgA isotypes across the vaccination time-course. Trend-lines display mean values per time point across all donors with available data; error-bars depict standard error of means.

**Supplementary Figure 4.**
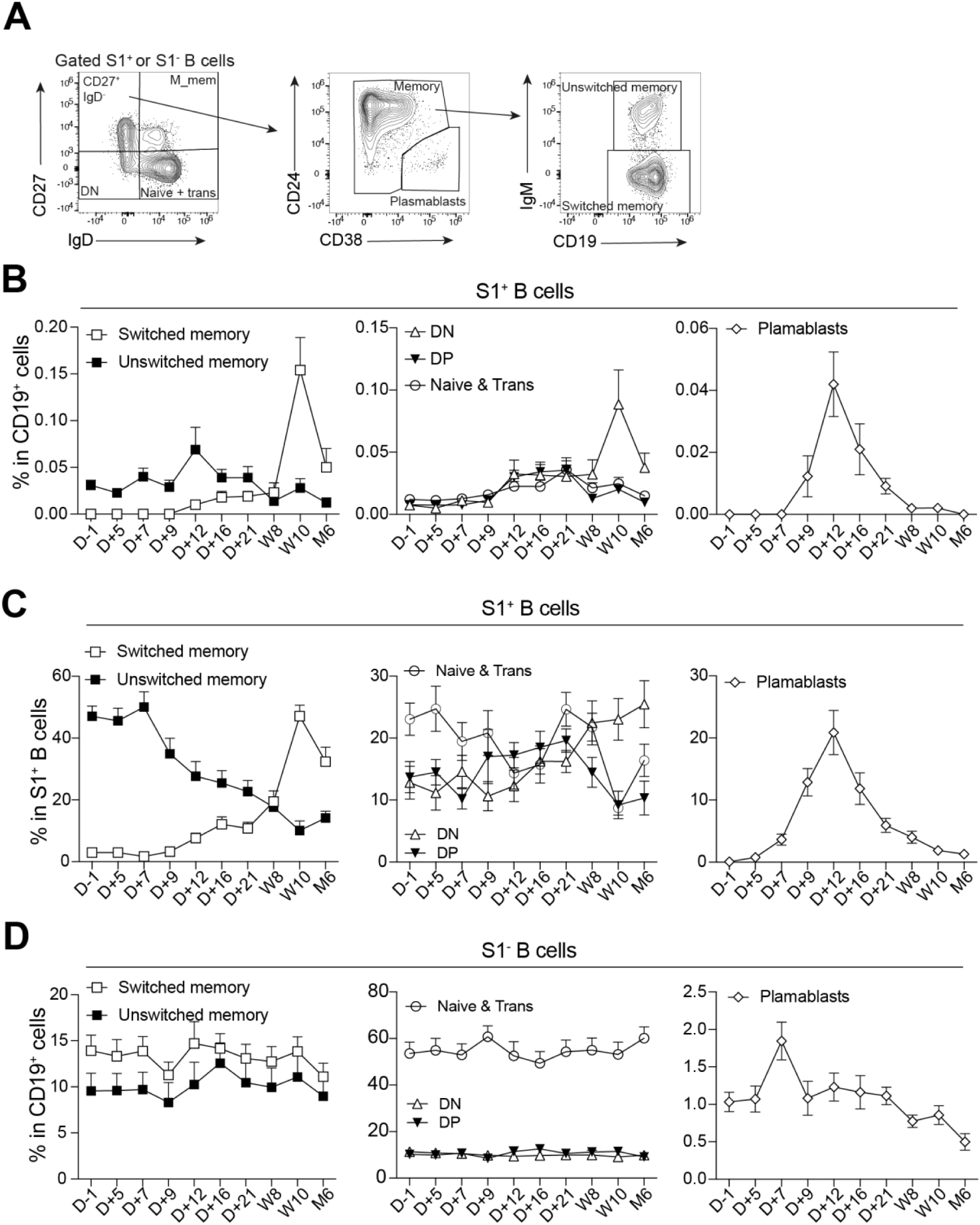
S1^+^ and S1^-^ B cell immunophenotyping. (A) Gating strategy for identification of vaccine-derived antigen-specific (S1^+^) or general B cell population (S1^-^) class-switched memory B cells (CD19^+^CD27^+^IgD-IgM^-^), unswitched memory B cells (CD19^+^CD27^+^IgD^-^IgM^+^), IgM memory (M_mem) B cells (CD19^+^CD27^+^IgD^+^), double-negative (DN) B cells (CD19^+^CD27^+^IgD^+^), naive plus transitional B cells (CD19^+^CD27^-^IgD^+^), and plasmablasts (CD19^+^CD27^+^IgD^-^CD24^-^CD38^+^). (B-C) Frequencies of S1^+^ class-switched memory B cells (empty squares), S1^+^ unswitched memory B cells (filled squares), S1^+^ DP B cells (invested filled triangles), S1^+^ DN B cells (empty triangles), S1^+^ naive plus transitional B cells (empty circles), and S1^+^ plasmablasts (empty diamonds) as (B) percentage of CD19^+^ cells and (C) percentage of S1^+^ B cells during vaccine response quantified using flow cytometry data. (D) Frequencies of S1^-^ class-switched memory B cells (empty squares), S1^-^ unswitched memory B cells (filled squares), S1^-^ DP B cells (invested filled triangles), S1^-^ DN B cells (empty triangles), S1^-^ naive plus transitional B cells (empty circles), and S1-plasmablasts (empty diamonds) as percentage of CD19^+^ cells during vaccine response quantified using flow cytometry data.

**Supplementary Figure 5.**
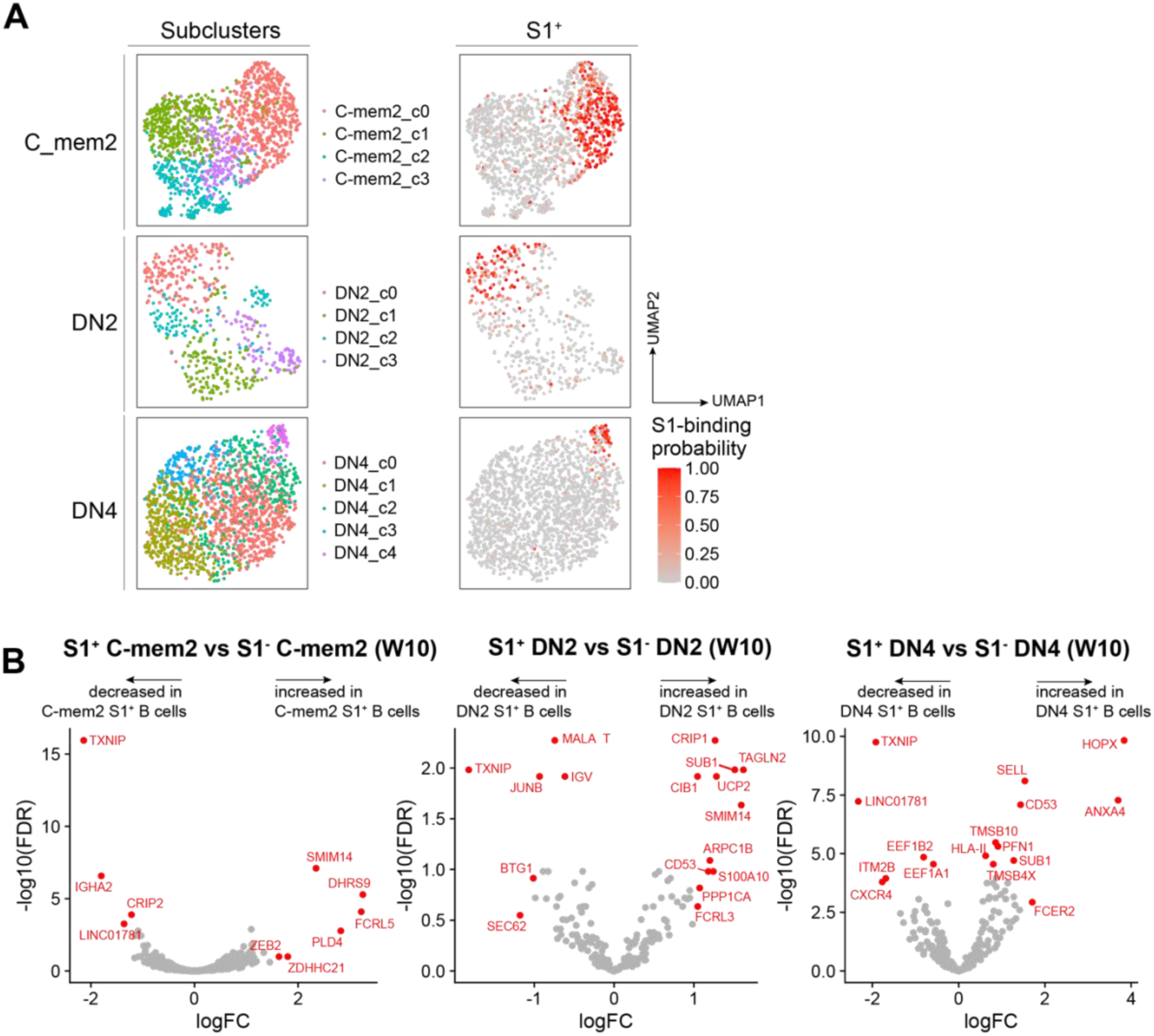
Gene expression signatures of S1^+^ B cells across B cell subsets. (A) Subclustering of C-mem2, DN2 and DN4 subsets and visualized in UMAP projections computed separately for each subset (left). S1^+^ B cells in these subclustered data were highlighted by their S1-binding probability score in the UMAP projection (right). (B) Differential gene expression analysis for genes of subclusters enriched in S1^+^ B cells compared to cluster not enriched in S1^+^ B cells at W10 for C_mem2, DN2 and DN4.

**Supplementary Figure 6.**
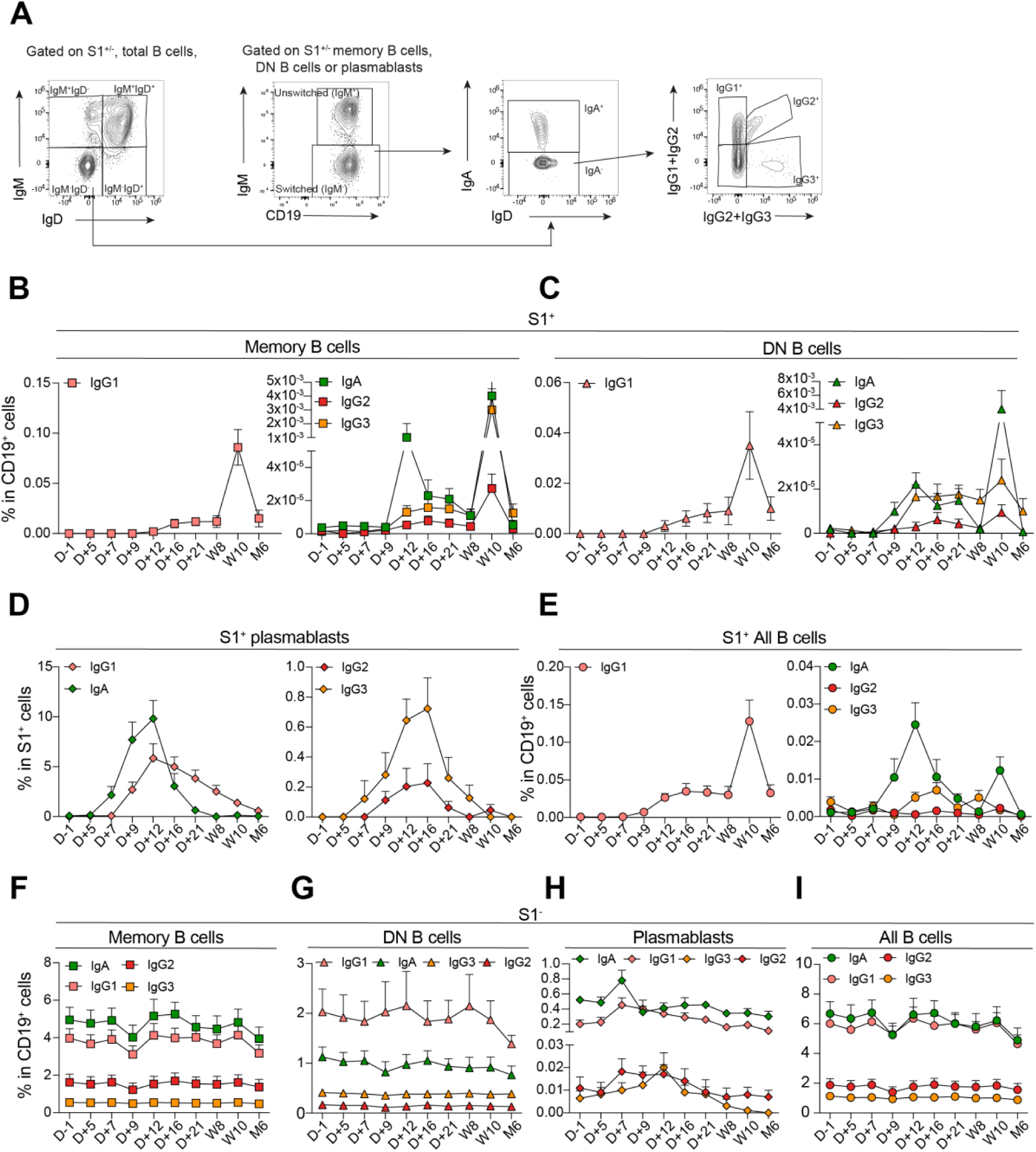
BCR isotype analysis in flow cytometry for S1^+^ and S1^-^ B cells. (A) Gating strategy for identification of IgA^+^, IgG1^+^, IgG2^+^ and IgG3^+^ class-switched total B cells (CD19^+^IgD^-^IgM^-^) class-switched memory B cells (CD19^+^CD27^+^CD24^+^CD38^lo^IgD^-^IgM^-^), class-switched double-negative (DN) B cells (CD19^+^CD27^-^IgD^-^IgM^-^), and class-switched plasmablasts (CD19^+^CD27^+^IgD^-^CD24^-^CD38^+^IgD^-^IgM^-^) in both vaccine-derived antigen-specific (S1^+^) or general B cell population (S1^-^). (B-E) Frequencies of S1^+^ class-switched memory B cells (CD19^+^CD27^+^IgD^-^IgM^-^, squares), class-switched double-negative (DN) B cells (CD19^+^CD27^-^IgD^-^IgM^-^, triangles), class-switched plasmablasts (CD19^+^CD27^+^IgD^-^CD24^-^CD38^+^IgD^-^ IgM^-^, diamonds) and class-switched total B cells (CD19^+^IgD^-^IgM^-^, circles) as percentage of CD19^+^ or S1^+^ B cells grouped by BCR isotype quantified using flow cytometry data during vaccines response. (F-I) Frequencies of S1^-^ class-switched memory B cells (CD19^+^CD27^+^IgD^-^IgM^-^, squares), class-switched double-negative (DN) B cells (CD19^+^CD27^-^IgD^-^IgM^-^, triangles), class-switched plasmablasts (CD19^+^CD27^+^IgD^-^CD24^-^CD38^+^IgD^-^ IgM^-^, diamonds) and class-switched total B cells (CD19^+^IgD^-^IgM^-^, circles) as percentage of CD19^+^ B cells grouped by BCR isotype quantified using flow cytometry data during vaccines response.

**Supplementary Figure 7.**
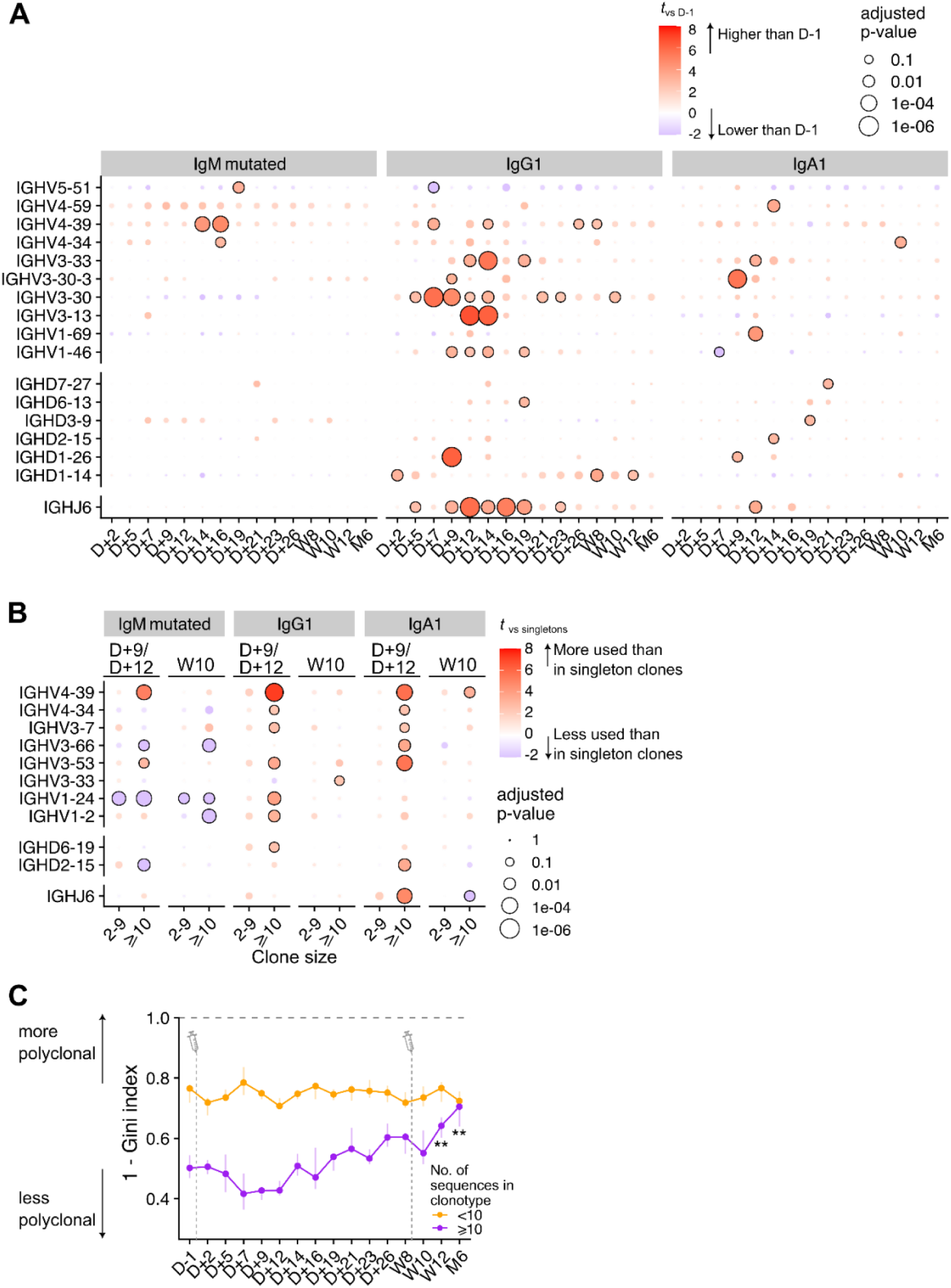
Transient polyclonal VDJ usage and clonal expansions in vaccine-induced primary response. (A) Change in immunoglobulin heavy-chain variable (V), diversity (D) and J (joining) gene usage across timepoints in the bulk BCR sequencing data. Gene usage was evaluated on subsets of BCR sequences which are mutated (<99% sequence identity to the germline) and of the IgM isotype (“IgM_mutated”), IgG1 and IgA1. Statistical significance was assessed by fitting mixed-effect linear models of percentage gene usage (dependent variable) against time point as the fixed effect and donor identifiers as the random effect. Bubble color depict effect size compared to D-1 (positive value indicates elevated usage of gene compared to D-1), and bubble sizes correspond to p-value after false-discovery rate adjustment. (B) Comparison of heavy-chain V, D and J gene usage between clonotypes of different sizes. Gene usage was computed for sequence subsets as defined in (A), separately for time points at the first peak (D+9 and D+12) and the second peak (Wk10). Clonotypes were grouped by their sizes, into singletons, clonotypes with 2-9 sequences and those with more than 10 sequences (“>=10”). Mixed effect models were fitted to compare gene usage against singleton clonotypes as a control, with clonotype sizes as fixed effects and donors as random effects. (C) Clonal diversity (as measured using the formula 1 – Gini index) of clonotypes with fewer than 10 sequences (<10) versus those with at least 10 sequences (>=10). Mixed effect models were fitted to compare against D-1 as control, using time points as fixed effects and donor as random effects. **, false-discovery rate < 0.01.

**Supplementary Figure 8.**
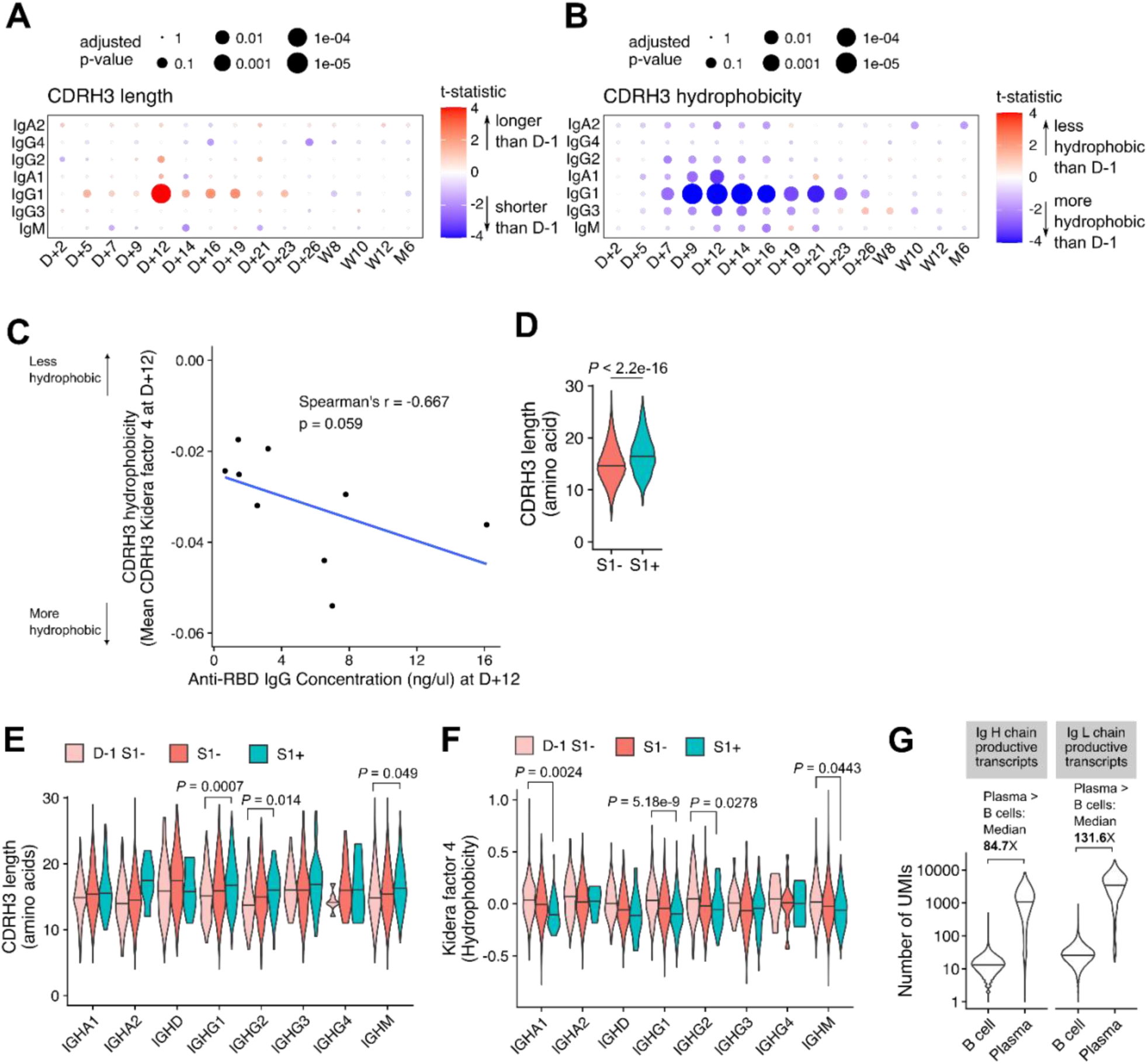
Longer and more hydrophobic CDRH3 is a feature of the vaccine-induced primary antibody response. (A-B) Comparison of CDRH3 amino acid sequence length and (B) hydrophobicity (using the Kidera factor 4 [Kidera et al. 1985] as a proxy) across time points. Sequences from the bulk BCR sequencing data were separated by isotypes and separate mixed effect models were fitted for CDRH3 length and Kidera factor 4 to compare each time point against D-1 as control, with donors as random effects. Bubble color indicates t-statistic from the mixed effect models, whilst bubble size indicates p-value after false-discovery rate correction. (C) Association between Anti-RBD IgG concentration as measured using ELISA (horizontal axis) and mean CDRH3 Kidera Factor 4 (indicating hydrophobicity) measured using D+12 sera. (D) CDRH3 amino acid sequence length distributions for S1^+^ and S1^-^ B cells. Statistical comparison was performed using a Wilcoxon rank-sum test. (E-F) Comparison of CDRH3 (E) amino acid length and (F) hydrophobicity (using the Kidera factor 4 [Kidera et al. 1985] as a proxy) between S1^+^ and S1^-^ B cells profiled in the scBCR-seq data. Distributions were visualized as violin plots separately for different BCR isotypes. S1^-^ data from D–1 was visualized separately to represent the baseline. Statistical comparisons of the D-1 S1^-^ baseline against S1^+^ were computed using Wilcoxon rank-sum tests and *p*-values were adjusted for multiple test corrections using the Benjamini-Hochberg method. (G) Comparison of the copy number (number of unique molecular identifiers [UMI] recorded per unique heavy-chain/light-chain productive transcript sequence) between B cells and plasma cells profiled in the scBCR-seq data.

**Supplementary table 1:** participants demographic details. CMV: human cytomegalovirus M:male, F:female.

| Participant ID | Sex | Age | CMV status |
| --- | --- | --- | --- |
| P01 | F | 34 | IgG Negative |
| P02 | F | 24 | IgG Positive |
| P03 | M | 32 | IgG Positive |
| P04 | M | 24 | IgG Positive |
| P05 | M | 28 | IgG Negative |
| P06 | F | 30 | IgG Negative |
| P07 | F | 29 | IgG Positive |
| P08 | F | 29 | IgG Positive |
| P09 | F | 28 | IgG Positive |
| P10 | F | 29 | IgG Positive |
| P11 | F | 29 | IgG Positive |
| P12 | F | 27 | IgG Negative |
| P13 | F | 29 | IgG Positive |
| P14 | M | 35 | IgG Positive |
| P15 | F | 24 | IgG Negative |
| Average/ratio | 73% F | 28.7 | 66% IgG positive |

